# SIZE DETERMINATION AND MULTIPLEXED FLUORESCENCE-BASED PHENOTYPING OF SINGLE CELL-DERIVED MEMBRANE VESICLES USING A NANOFLUIDIC DEVICE

**DOI:** 10.64898/2026.04.17.719178

**Authors:** Quentin Lubart, Sune Levin, Viktoria de Carvalho, Elin Persson, Stephan Block, Silver Jõemetsa, Erik Olsén, Sriram KK, André Görgens, Samir El-Andaloussi, Fredrik Höök, Marta Bally, Fredrik Westerlund, Elin K. Esbjörner

## Abstract

Extracellular vesicles (EVs) are cell-secreted biological nanoparticles that play a crucial role in intercellular communication and are gaining increasing attention as diagnostic biomarkers, therapeutic agents, and drug delivery vehicles. Consequently, the development of robust and sensitive methods for their characterization is essential. Herein we present the use of a microscope-mounted nanofluidic device for direct size determination and multi-parametric (3-color) fluorescence-based phenotyping of single biological nanoparticles that are in the size range of 20-200 nm in a method we denote Nano-SMF (SMF; size and multiplexed fluorescence). We demonstrate that it is possible to accurately determine the size of nanoparticles by analyzing their one-dimensional Brownian motion during directional flow through nanochannels, achieving size distributions for monodisperse nanoparticle solutions that are on par with TEM analysis, and size discrimination of nanoparticle mixtures that is significantly improved compared to conventional nanoparticle tracking analysis (NTA). Furter, we demonstrate that the method can be applied to analyze EVs directly in minute volumes of cell supernatant, avoiding pre-isolation or concentration steps. The method was applied to phenotype CD63- and CD81-positive EVs from a human embryonic kidney cell model, demonstrating that vesicle sub-populations defined by these two tetraspanin biomarkers differ significantly in size.

## INTRODUCTION

Cells release a variety of compositionally diverse membrane-enclosed vesicles into their environment.^1,2^ These entities, collectively referred to as extracellular vesicles (EVs), have the capacity to mediate intercellular communication and transduce signals via the transfer of biological content such as proteins, lipids, metabolites, and nucleic acids, to recipient cells.^3^ EVs regulate a diverse range of normal physiological processes, but can also have significant pathophysiological roles,^4^ highlighting their potential as diagnostic biomarkers and putative drug targets.^5^ EVs are also being exploited as therapeutics, for example as effectors in regenerative medicine or as drug delivery vehicles.^6–10^ Precise characterization of EVs, and particularly the mapping of EV sub-populations, is therefore crucial but remains difficult to achieve with existing methods.^11,12^

EVs have a wide size variation that ranges from the nano- to micrometer regime.^13^ Challenges in characterization of small EVs (< 200 nm) include their size, transparency, significant heterogeneity with existence of overlapping sub-populations,^14–16^ and relatively low concentrations in culture media and bio-fluids.^13^ This makes them hard to isolate into pure sub-fractions.^17^ Most EV isolation methods rely on initial separation and enrichment protocols, followed by some type of ensemble averaging identification of specific protein markers and/or cargoes in the enriched fractions, using for example Western blotting, enzyme-linked immunosorbent assays, mass spectrometry, or bead-coupled flow cytometry.^13,18,19^ EV size distributions have typically been determined using electron microscopy (EM) or nanoparticle tracking analysis (NTA) commonly used in dark field scattering mode.^20^ However, neither of these methods are ideal for multiplexed analysis and while fluorescence-based detection is in principle possible in NTA, many commercial instruments are often limited to one color,^21^ and those that enable multi-color colocalization usually do not provide coupled measurements of size for the colocalized particles. Although several interesting developments have been made to enable multiplexed phenotyping of EVs, including various adaptations of conventional flow cytometry instruments,^22^ development of high sensitivity flow cytometry,^23^ and application of various optical sensing platforms,^24–31^ it remains highly challenging to simultaneously record size and several phenotypical fingerprints of individual EVs. Furthermore, most single particle methods have severely limited capabilities to accurately detect and characterize particles smaller than 50 nm, and to distinguish particle sub-populations in the sub-100 nm range. This limits the analysis of EVs, particularly sub-populations that derive from multi-vesicular bodies, commonly referred to as exosomes,^2^ but also of other types of small biological particles such as synaptic vesicles which are typically ∼40 nm in diameter.^32^

We have previously demonstrated that it is possible to use nanochannel devices, originally developed to visualize single large DNA molecules,^33,34^ to image and count single fluorescent biological nanoparticles on a standard epi-fluorescence microscope.^35^ This enabled accurate particle concentration determination over a relatively broad interval, as well as colocalization of two colors, but without size determination. In parallel, we devised a two-dimensional flow nanometry method that combines accurate quantification of nanoparticle size, based on lateral diffusion, and intensity (scattering/fluorescence), but only if the object of interest is tethered to a laterally mobile supported lipid bilayer.^36^ Herein, we combine these two developments into a method for combined size determination down to 20 nm, and simultaneous multiplexed fluorescence-based phenotyping with up to three individual colors for single biological nanoparticles in suspension. The size determination is enabled by tracking the one-dimensional Brownian motion of the particles in the nanochannels and allows for detection of thousands of individual particles per minute. We apply the method, which we denote Nano-SMF (SMF; size and multiplexed fluorescence), to detailed phenotyping of EVs isolated from a human embryonic kidney cell line, showing that sub-populations of EVs positive for the common EV-associated tetraspanins CD63 and CD81 differ substantially both with respect to size and tetraspanin density in their membrane. Additionally, we demonstrate that Nano-SMF can be used to directly analyze EVs in minute volumes of conditioned cell media without any pre-isolation and concentration steps. We believe that this new method holds significant potential as a complement to existing ensemble-averaging biochemistry-based technologies and could provide new information and characterization, not only to cell-derived vesicles, but also to other types of biologically relevant soft nanoparticles such as synaptic vesicles, viruses, and drug delivery vehicles, such as lipid nanoparticles.

## RESULTS AND DISCUSSION

### Size determination of nanoparticles in flow

The Nano-SMF method is based on confinement of EVs to a nanofluidic device (Fig. 1a) which is mounted on an inverted epi-fluorescence microscope (Fig. S1), allowing single nanoparticles to be imaged in real-time and in solution, as they flow through an array of parallel nanochannels with 300x300 nm^2^ cross section (Fig. 1b).^35^ Using nitrogen pressurized flow, we achieved flow rates that allowed us to determine nanoparticle size based on their Brownian motion in the channels, and we demonstrate that it is possible to collect at least three different fluorescent signals per particle simultaneously.

**FIGURE 1:**
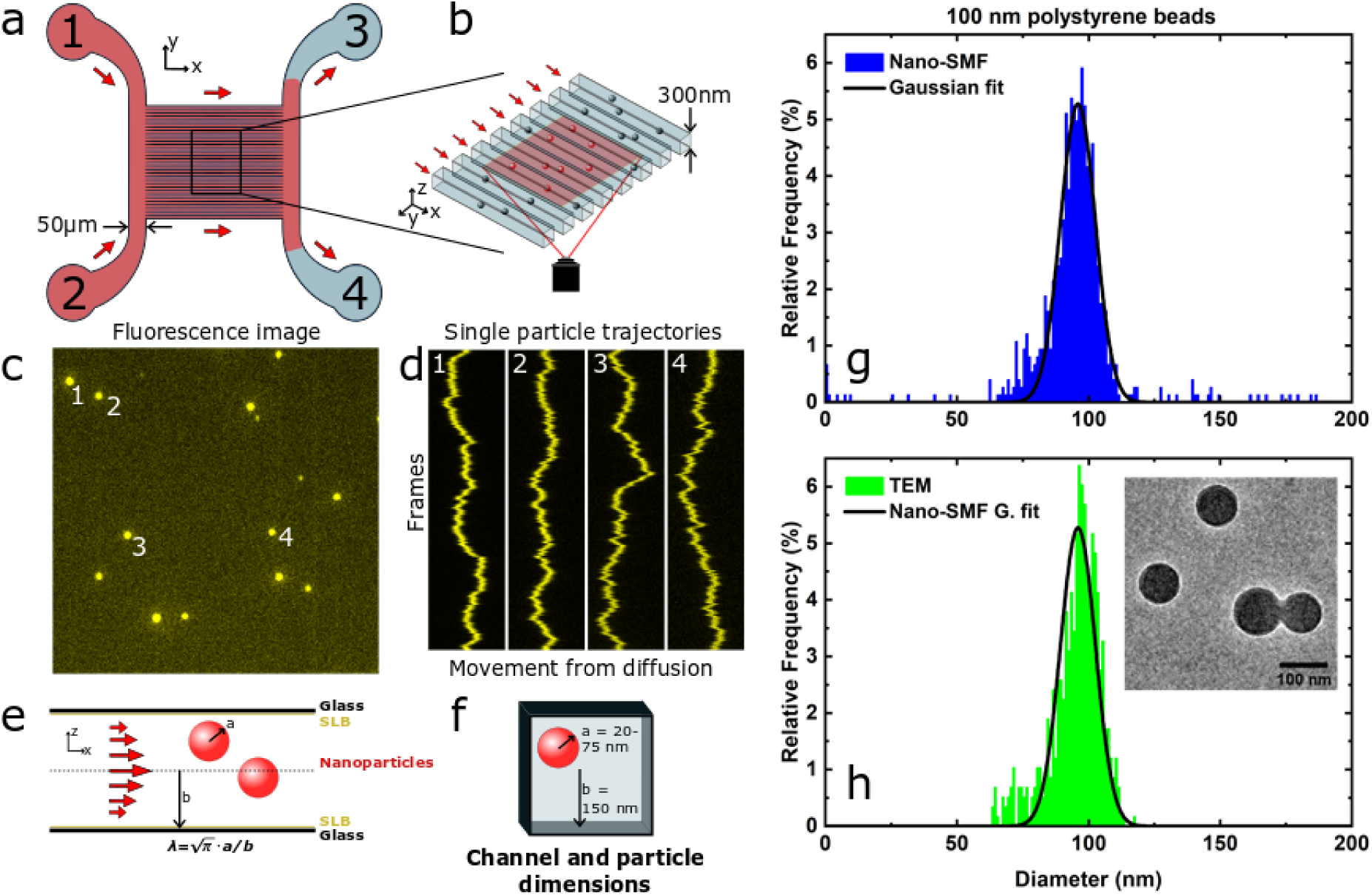
Summary of Nano-SMF methodology and output. a) Schematic of the nanofluidic chip. b) Representation of nanoparticles flowing through the nanochannels on top of an epifluorescence microscope. c) Fluorescence image of 100 nm polystyrene beads in nanochannels. d) Tracking of representative particles through a video with numbers from c). e) Representation of nanoparticles inside a nanochannel coated with a supported lipid bilayer (SLB). f) Schematic of nanoparticle and nanochannel dimensions. g) Size distribution of 100 nm polystyrene beads determined using Nano-SMF, fitted with a Gaussian distribution. h) Size distribution of 100 nm polystyrene beads determined using transmission electron microscopy compared with the Gaussian distribution detected with Nano-SMF. Inset displays a snapshot of the polystyrene beads imaged with transmission electron microscopy.

First, we recorded videos of fluorescent 100 nm polystyrene beads flowing through the nanochannels (Fig. 1c; Supplementary Video 1), demonstrating that the nanochannel confinement enables true capture of particles one-by-one. This avoids problems with swarm detection (multiple vesicles simultaneously detected as one unit), which are common in conventional flow cytometers operating in small particle mode.^37^ A particle tracking algorithm was developed to determine the individual trajectory of each particle (Fig. 1d, Supplementary Note 1). The accuracy and sensitivity of the detection was greatly facilitated by the nanochannels keeping each particle moving on a single track and in focus throughout the imaging experiment. The motion of each tracked particles was decomposed into stochastic (diffusion) and deterministic (drift in flow direction) components (Supplementary Note 2) as previously described for nanoparticles tethered to a bilayer surface.^38^ The stochastic component entails information about hydrodynamic size of each particle. Using this approach to derive diffusion coefficients, we calculated the hydrodynamic diameter of each individual nanoparticle using the Stokes–Einstein relation, with corrections accounting for the hydrodynamic coupling between the nanoparticles and the nanochannel walls (Fig. 1e). The correction was implemented through a hindrance factor ***H***, which depends on the ratio of the nanoparticle radius ***r*** to the half nanochannel side length ***a*** (Fig. 1e and 1f) and was derived using the center line approximation for squared cross-section channels (see Supplementary Note 2). This implementation improved the accuracy in size determination compared to previously published equations describing the hydrodynamic coupling to the walls of nanopores or nanoslits^39^ (Fig. S2). The size determination accuracy of the Nano-SMF method will depend on particle flow rates and the ability to precisely determine particle positions through the imaging-based detection. We therefore optimized the setup by systematic variation of flow rates and camera exposure times (Fig. S3), showing that at a flow rate of 15 μm/s (corresponding to 10 mbar flow pressure), and an exposure time of 30 ms, we could record data with good signal-to noise and accurate read-out of particle size, whilst retaining the advantage of high throughput acquisition (detecting ca 200 particles per minute).

Using the optimized Nano-SMF setup, we determined the mean size of the fluorescent polystyrene beads to 95.9 ± 6.6 nm (Fig. 1g), which was in excellent agreement with TEM-based analysis (97.7 ± 6.9 nm; Fig. 1h) as well as the nominal size provided by the manufacturer (100 ± 10 nm). Furthermore, using flow rates and particle counts (Supplementary Note 4), we determined the concentration of the polystyrene particle solution to 2.2•10^10^ p/mL in good agreement with the information provided by the manufacturer (4•10^10^ p/mL) and well within the error range for particle concentrations reported in a previous study.^35^

An important feature in methods for single fluorescent nanoparticle analysis is the ability to accurately detect and characterize particles even when variations in their sizes and intensities are large. In most intensity-reliant analytical methods, variation in particle intensity leads to contrast problems. This is exemplified by the snapshot images in Fig. 2a and kymographs in Fig. 2b showing a sample consisting of a mixture of 50 and 100 nm sized fluorescent polystyrene beads with an average emission intensity difference of 10.4 (insert in Fig. 2e). Nano-SMF can overcome this problem by accurately recapitulating the size distribution of the two particle types, both separately (Fig. 2c and 2g) and, more importantly, as a mixture (Fig. 2e). By contrast, a conventional nanoparticle tracking analysis (NTA) instrument, operating in scattering mode, could not detect the 50 nm polystyrene beads (Fig. 2f), presumably due to the significant differences between the scattering intensities of the particles (*I ∝ r*^6^, inset Fig. 2e). The nanochannel confinement and parallel arrangement separates the nanoparticles in space and simultaneously keeps them in focus throughout the measurement. This prevents brighter particles from overshadowing dimmer ones and their aligned flow along the nanochannels furthermore hinders crossing of particle trajectories which allows long tracking times (≈ 400 frames). As a result, the measured size distributions for single sized polystyrene beads were significantly narrower in Nano-SMF compared to NTA (Fig. 2d, 2h) and the mean was identified with higher precision (95.9 ± 6.6 nm in Nano-SMF compared to 87.3 ± 25.3 nm in NTA). This comparison shows that the Nano-SMF method offers greater dynamic range as well as better precision and accuracy in size determination compared to NTA.

**FIGURE 2:**
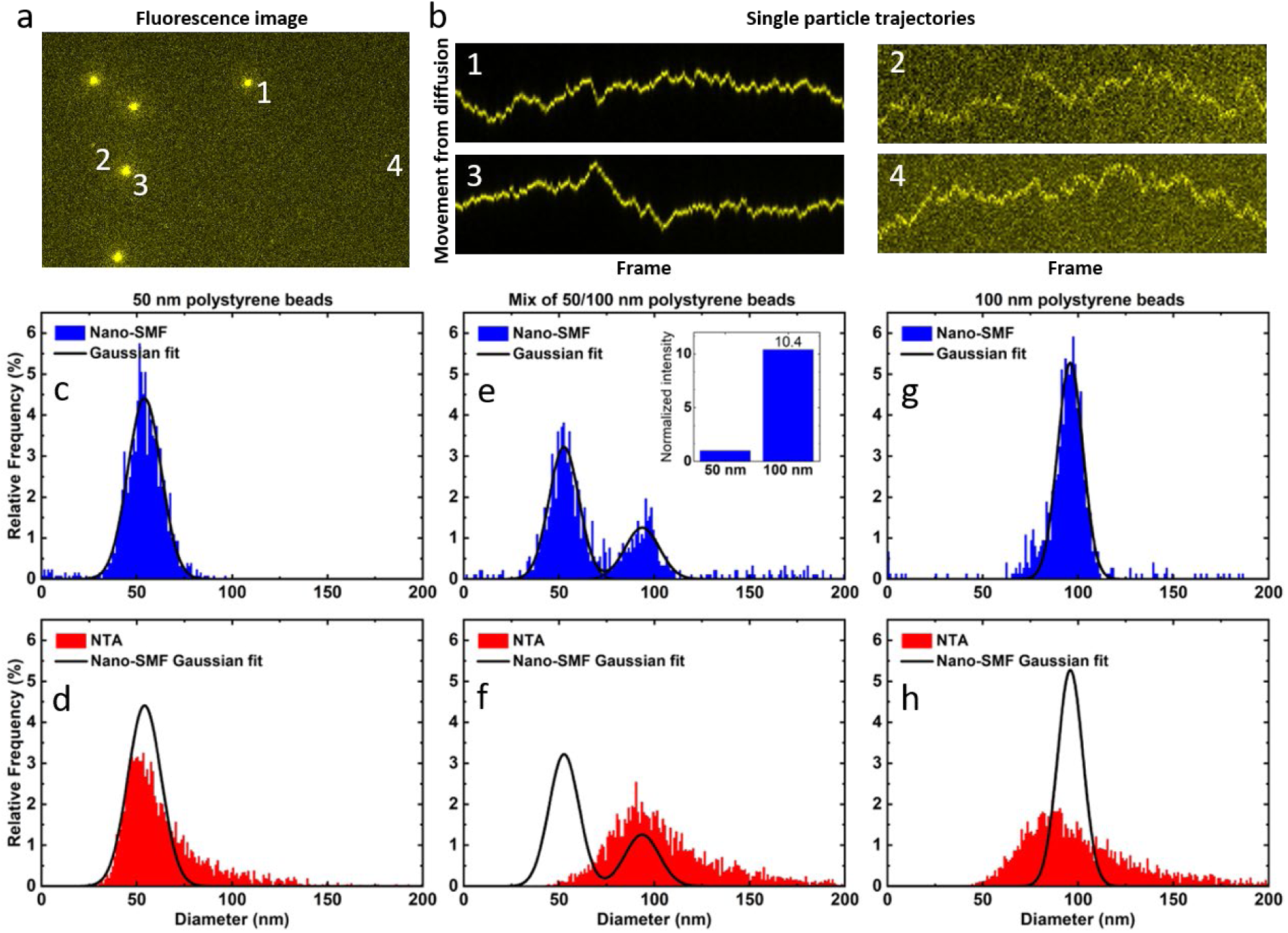
Comparison of size distributions for 50 and 100 nm polystyrene beads obtained using Nano-SMF (blue) and NTA (red). a) Snapshot of mixed 50 and 100 nm polystyrene beads using Nano-SMF. b) Trajectories displaying Brownian motion from numbered particles in (a). c-d) Size distribution of 50 nm polystyrene beads obtained with Nano-SMF (c) and NTA (d). e-f) Size distribution of mixed 50 and 100 nm polystyrene beads obtained with Nano-SMF (e) (insert displays the difference in fluorescence intensity between the 50 and 100 nm particles) and NTA (f). g-h) Size distribution of 100 nm polystyrene beads obtained with Nano-SMF (g) and NTA (h). Size distributions obtained with Nano-SMF (c, e and g) are fitted with Gaussian distributions and displayed for comparison with NTA (d, f and h).

Next, we tested the lower limit for the size detection (which is often restricted by particle brightness) by analyzing fluorescent polymeric nanoparticles (StarBright Violet 670-labelled P-dots) with a nominal diameter of 20 nm, recapitulating their reported mean size and narrow size distribution (20.2 ± 4.7 nm, Fig. S4) without any apparent cut-off on the low diameter side of the histogram. This suggests that Nano-SMF pushes the lower limit of detection further than that of other nanoparticle sizing platforms.^23–25,40–45^

The nanochannel confinement method presented here is not restricted to fluorescence-based detection. It has been shown that single particles, down to the size of proteins, can be detected in single nanochannels using dark field scattering microscopy.^46^ We tested if this concept could be extended to the nanochannel setup used here, and to using the dark field scattering mode on a standard fluorescence microscope, by analyzing gold nanoparticles with nominal diameters of 60 nm and 100 nm. The particles could be well detected, and their sizes could be recapitulated with the same precision as by TEM (Fig. S5).

### Simultaneous size determination and co-localization of multiple fluorescent signals

Multiplexing is important for versatile single particle characterization. We therefore explored the capacity of the Nano-SMF setup with respect to simultaneously probing the size and multiple fluorescent signals of single large unilamellar membrane vesicles (LUVs). Fig. 3a shows the size distribution of LUVs composed of POPC lipids and 2 mol% rhodamine-labelled DHPE. These LUVs had been sequentially extruded through filters with 100 nm and 30 nm pore size to achieve an average LUV size below 100 nm to resemble size distributions of extracellular vesicles (EVs). The mean size of the LUVs was determined to 66 ± 30 nm whereas NTA yielded a mean size of 85 ± 28 nm (Fig. S6, Table S1), possibly due to the challenge in NTA of accurately sizing particles below 50 nm.^20,47^ The fluorescence intensity of individual LUVs correlated strongly with their size (Fig. 3b) and the scaling exponent (𝐼𝐼 ∝ 𝑟𝑟^𝑋𝑋^) was X=2.54 ± 0.02 which is slightly higher than the expected scaling with the surface area of a sphere (X = 2), but in good agreement with published data^48,49^ and consistent with the observation that a fraction of extruded LUVs typically are multilamellar.^49^ As an alternative, we labelled POPC LUVs with the lipophilic dye PKHred, as this is a common method for post-labelling of EVs and other biological membrane-enclosed particles.^50,51^ The PKHred-labelled POPC LUVs had a mean size of 68 ± 19 nm according to our Nano-SMF method, compared to a mean size of 77 ± 24 nm determined by NTA (Fig. S7a, Table S1). When we evaluated the relationship between fluorescence intensity and size (Fig. S7b) we obtained a correlation with a scaling exponent of *X* = 1.15 ± 0.02, which is lower than the theoretical expected value (*X* = 2), but in good agreement with recently reported data.^48^ A possible reason to this might be micelle formation of the PKHred dye, which is a commonly reported problem for PHK dyes,^52^ which possibly can be discerned as an additional population in Fig. S7b, skewing the regression line to a flatter slope.

**FIGURE 3:**
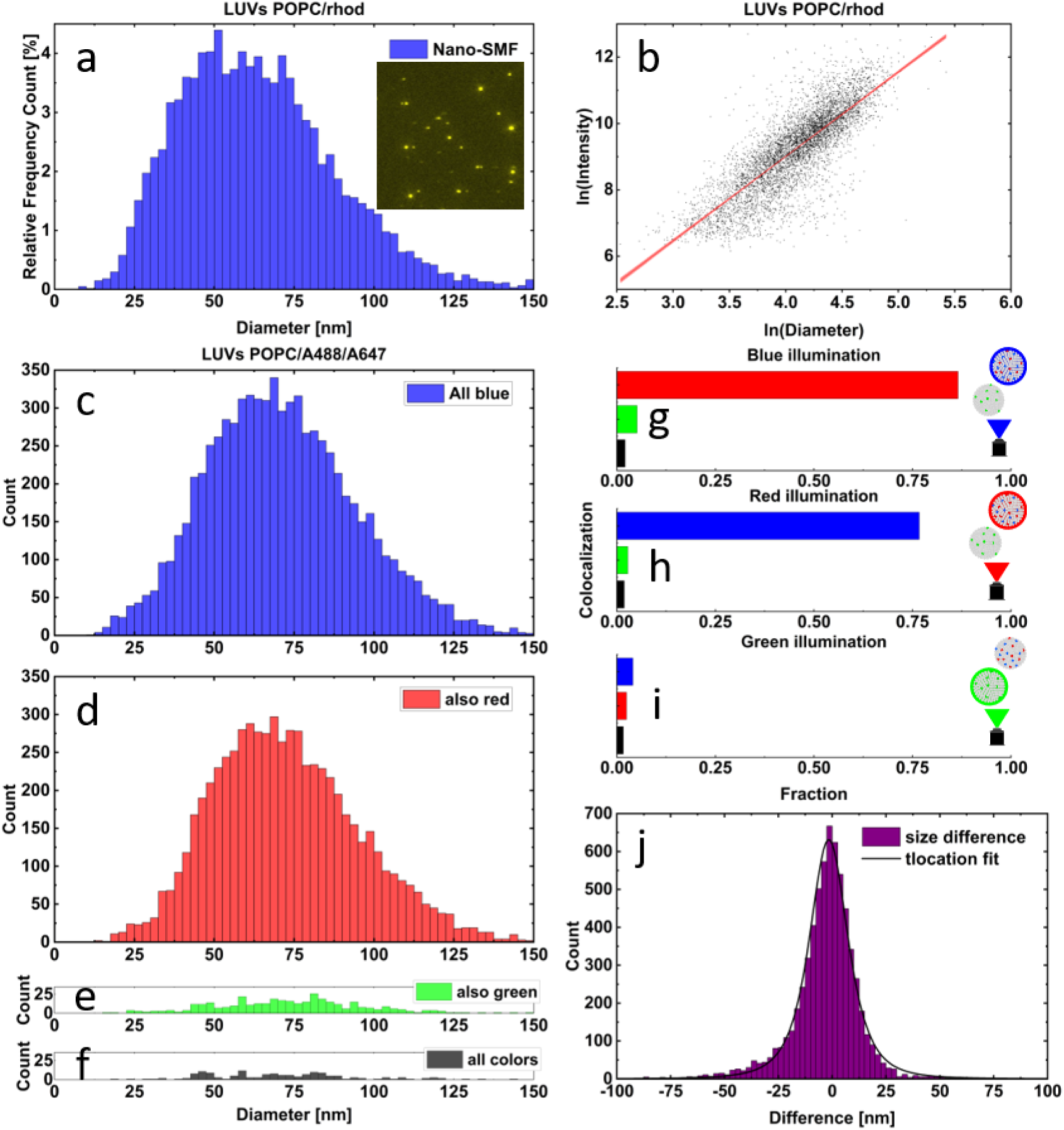
Demonstration of colocalization with the Nano-SMF for a sample of POPC LUVs. a) Size histogram obtained using Nano-SMF on LUVs POPC/rhod. Inset displays a snapshot of the LUVs. b) Logarithmic scatter plot of size vs intensity for LUVs POPC/rhod. The linear fit gives a value of 2.54 for the size-intensity scaling. c-f) Colocalizing size histograms for all particles collected with blue illumination on LUVs POPC/A488/A647. Histogram of all detected particles (c) or particles colocalizing between blue and red (d), between blue and green (e) or between all three color channels (f). g-i) Fractions of colocalized particles depending on the wavelength of illumination. A large fraction of the particles detected in blue illumination displays colocalization with the particles detected with red illumination (g), and vice versa (h). A low fraction of the particles detected with green illumination (i) colocalize with particles detected in the other color channels. j) Size difference between colocalized particles with sizes determined in both the blue and red channels, t-location fit parameters σ = 9.5 and peak location μ = 2.8.

One significant advantage with the Nano-SMF method is the possibility to track single nanoparticles for extended time periods, which should enable straightforward multi-color intensity acquisition by sequential fast switching of the excitation source. To test this, we prepared two types of POPC LUVs; one containing both 2% ATTO488 and 2% ATTO647 labelled lipids and one labeled with 2 % Rhodamine B (excitation max at 546 nm) labelled lipid. We mixed the two LUVs and imaged them in three color channels (excitation at 475, 555 and 630 nm, hereafter referred to as blue, green and red respectively, with 20.3 fps; 6.8 fps per color) using a multi-bandpass filter (Fig. S8) to collect the emission. Fig. 3c shows the size distribution histogram for all LUVs detected with blue excitation, after correction for cross-excitation (Supplementary Note 3). Fig. 3d shows the corresponding size distribution for the population of particles in Fig. 3c that were also detected with red excitation (data corrected for cross-excitation). Fig. 3g shows the proportion of blue particles co-detected with red, green, or all three excitation wavelengths, where almost all particles detected with blue excitation are also detected with red excitation, and very few are detected with green excitation. These findings show that Nano-SMF captures, with high fidelity, the colocalization of the ATTO488 and ATTO647 dyes to one of the LUV types in the mixed sample. A similar high degree of colocalization was observed by analyzing the fraction of LUVs detected by red excitation that were also excited by blue excitation (Fig. 3h). Conversely, the fraction of liposomes in Fig. 3c that were also detected with green excitation (Fig. 3e) was low (5.1%, Fig. 3g); and a similar value (2.7%) was obtained when probing colocalization by red and green excitation (Fig. 3f, 3h) or probing for the fraction of liposomes detected by green excitation that were detectable by both the other excitations (4.0% and 2.4% for blue and red respectively, Fig. 3i). This demonstrates that Nano-SMF analysis clearly distinguishes the Rhodamine-labelled LUVs from the dual labelled ATTO488/ATT647 ones, and that Nano-SMF has a low false detection rate. Furthermore, the size distribution of the LUVs is independent of the excitation wavelength used for detection; the mean diameter of the ATTO488/ATTO647 LUVs was 71 ± 24 nm using blue excitation or 74 ± 24 nm using red excitation. A t-location scale distribution analysis of the size difference estimated following blue or red excitation of individual ATTO488/647 LUVs is shown in Fig. 3j and reveals a low error between the calculated size of the same particles using different excitations with standard deviation σ = 9.5 and peak location μ = 2.8. Altogether, the LUV experiments show that Nano-SMF can be used to determine both hydrodynamic size and perform multiplexed color colocalization with discrimination between different fluorescent signals with high accuracy, enabling the detection and characterization of sub-populations within a complex biological sample. This data was compared with NTA in scattering (and fluorescence mode when possible) (Table S2) showing a higher overall mean size (≈10-20 nm) and an overestimation of 10 nm in fluorescence mode compared to scattering mode.

### Phenotyping of Extracellular Vesicles (EVs)

Next, we applied Nano-SMF to analyze EVs derived from human embryonic kidney (HEK) cell cultures. To challenge capacity of Nano-SMF in distinguishing EV sub-populations based on their size and molecular identity, the cultures were grown with a mixture of genetically engineered and wild-type (wt) cells. The engineered cells had been modified by lentiviral transduction to stably overexpress the tetraspanin CD63 fused to the fluorescent protein mNeonGreen (hereafter referred to as mNeon) on the intraluminal side;^53^ CD63 is highly enriched in the intraluminal vesicles of multivesicular bodies, involved in exosome biogenesis,^54^ and thus widely used as a marker to track, label, and phenotype exosomes. ^18,55^ Figure 4a shows the size distribution of CD63-mNeon positive EVs with a mean size of 90 ± 40 nm. The size distribution deviates clearly from the Gaussian form observed for the polystyrene beads in Fig. 1g and for the LUVs in Fig. 3. This and subsequent EV distributions were instead fitted to a lognormal function (Table S3) to account for the subpopulation of larger particles. We also analyzed the relationship between the size and the CD63-mNeon abundance (intensity) (Fig. 4b). Notably, this data has considerably higher intensity variation compared to the synthetic LUVs (Fig. 3b). This suggests that the incorporation of CD63 into EVs is not as homogenous as the distribution of dye-labelled lipids into a POPC LUV during dry film hydration. Comparison of the size distributions of CD63 positive EVs to those of polystyrene beads and LUVs shows that LUVs and EVs display much higher degrees of size heterogeneity compared to beads (Fig. 4c), as also indicated by the calculated polydispersity index with Nano-SMF (Table S1 and Table S3, PDI_polystyrene beads_ = 0.005). This is likely a reflection of their biogenesis and related to the biological and physical processes regulating intraluminal vesicle formation inside multivesicular bodies. In addition, the size distribution of the CD63-mNeon EVs is significantly right-skewed as previously noted, modelled and proposed to be related to the dynamic scaling of domain growth on the limiting membrane of multivesicular bodies.^56^

**FIGURE 4:**
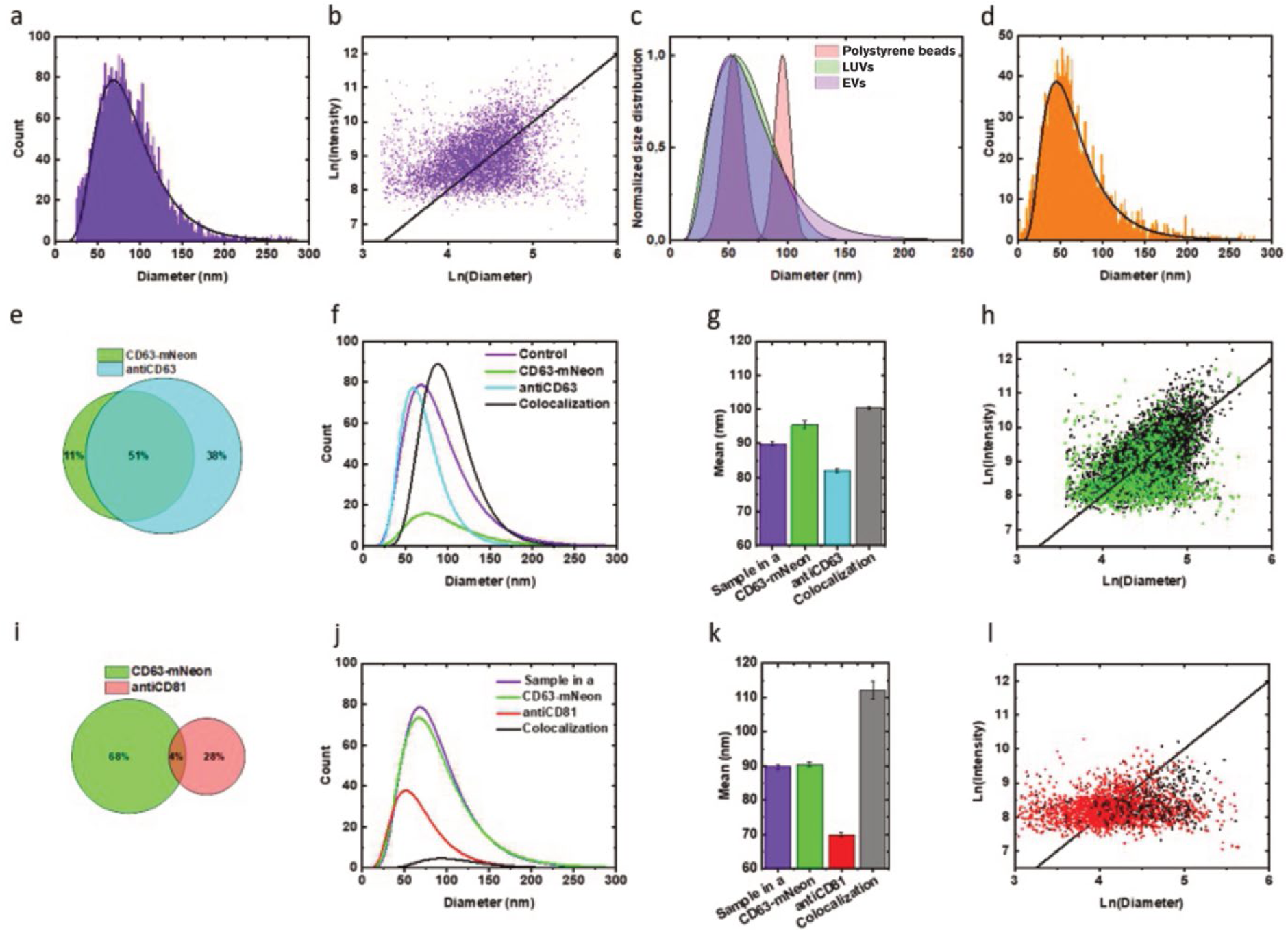
Comparison of size distributions and fluorescence-based phenotyping of biomarkers on exosomes using the Nano-SMF method. a) Histogram of CD63-mNeon exosomes and its logNormal fit used to determine mean hydrodynamic diameter and distribution. b) Fluorescence intensity versus hydrodynamic diameter ln-ln plot for the CD63-mNeon exosomes in (a). The line represents the theoretical slope of 2. c) Comparison of normalized fitted size distributions for 50 and 100 nm polystyrene beads (red), LUVs (POPC/RhoB) (green) and CD63-mNeon exosomes (blue). d) Histogram of non-purified, serum fresh, CD63-mCherry exosomes and its lognormal fit. e-h) Subpopulation characterization of CD63-mNeon exosomes (green) labelled with antiCD63 (light blue), the unlabeled CD63-mNeon exosomes in (a) are included as reference (violet). The particles colocalizing the CD63-mNeon and anti-CD63 markers are represented in black. e) Venn diagram of particle subpopulations distribution, the colocalization is represented by the overlapping region. f) Comparison of lognormal fitted size distributions. g) Comparison of the mean hydrodynamic diameters. h) Fluorescence intensity versus hydrodynamic diameter ln-ln plot, showing CD63-mNeon positive particles (green) and CD63-mNeon/anti-CD63 co-localizing particles (black). The line represents the theoretical slope of 2. i-l) Subpopulation characterization of CD63-mNeon exosomes (green) labelled with antiCD81 (red). The unlabeled CD63-mNeon exosomes in (a) are included as reference (violet). The colocalizing particles are represented in black. i) Venn diagram of particle subpopulations distribution, the colocalization is represented by the overlapping region. j) Comparison of lognormal fitted size distributions. k) Comparison of the mean hydrodynamic diameters. l) Fluorescence intensity versus hydrodynamic diameter ln-ln plot showing anti-CD81 positive particles (red) and CD63-mNeon/anti-CD81 co-localizing particles (black). The line represents the theoretical slope of 2.

Next, we labelled the HEK cell derived EVs with a fluorescent (APC) conjugated primary antibody reactive against human CD63 (anti-CD63) and recorded particle sizes and intensities of mNeon and APC simultaneously. Out of ∼12000 detected fluorescent particles (Table S3), 51% were positive for both CD63-mNeon and the anti-CD63 (sub-population denoted as colocalizing), whereas 38% stained with anti-CD63 only (Fig. 4d). This reflects the fact that the EVs in the sample originated from a mixture of CD63-mNeon expressing and wt cells and demonstrates that we can clearly distinguish engineered and wt EVs as two discrete populations, which, as we will discuss further below, also differ in size. We also identified a small fraction (11%) of CD63-mNeon positive particles that were unable to bind anti-CD63 (Fig. 4e-f; Fig. S15; Table S3), but we excluded them from further analysis since it is unlikely that they present vesicles with correctly folded and membrane spanning CD63, making their biological functionality questionable. The size distributions of the different detected sub-populations are shown in Fig. 4f (log-normal fit) and Fig. S15, and the corresponding mean diameters are shown in Fig. 4g and Table S3. Several interesting observations can be made from this experiment. First, by comparing the size of the CD63 mNeon-positive EVs analyzed in absence of antibody (Fig. 4a; violet line/bar in Fig. 4f, Fig 4g) to the sub-population that colocalizes with anti-CD63 (e.g. are antibody labelled), we observe an increase in hydrodynamic diameter of ∼11 ± 1 nm. This corresponds to an added protein layer on the surface of the EVs, due to antibody binding, with a thickness of ∼5.5 nm. This can be compared to the molecular dimensions of an IgG antibody of 14.5 nm x 8.5 nm x 4.0 nm^57^ (the longest dimension being between the tips of the Fab arms which bind the EV surface) and suggests that the antibody mainly binds parallel to the EV surface. The significant size increase would suggest that the EV surface is well coated with antibodies and thus that CD63-mNeon is highly abundant.^58^ This comparison demonstrates that Nano-SMF can be used, not only to determine the size of single biological nanoparticles, but also to quantify size changes resulting from the binding of macromolecules to their surfaces.

Moreover, it provides a useful estimate of how much the sizes of EVs are overestimated by the common use of antibodies for phenotypic detection. Second, our data shows that native anti-CD63 positive EVs are on average 82 ± 42 nm in size, including the antibody layer. Upon the reasonable assumption that the antibody layer thickness is independent of EV size, our data therefore suggest that CD63-positive EVs released from HEK cells are ∼71nm in diameter. This is slightly larger than reported in literature (∼ 55nm)^23^ and could be explained by that the EVs are coming from a different cell line and using a different protocol of isolation. A third observation is that overexpression of CD63-mNeon in the HEK cells increases the sizes of the released CD63-positive EVs by ∼18 nm (or 26%) to ∼89 nm. This is in good agreement with results obtained by fluorescence correlation spectroscopy-based size analysis, showing that CD63-GFP positive EVs from HEK293T cells have a mean size of ∼100 nm^59^ and indicates, furthermore, that the availability of CD63 in MVBs regulates vesicle size during exosome biogenesis. Fig. S16 reveals an expected good correlation between antibody binding density and presence of CD63-mNeon moieties on the EV surfaces. In addition, wt EVs bind, on average, significantly less anti-CD63 compared to CD63-mNeon positive ones as shown by the intensity comparisons in Fig. 4h. This higher abundance of CD63 in the EVs derived from producer cells with CD63-mNeon overexpression is consistent with previous observations;^22^ our results extends this view by suggesting that the abundance of this tetraspanin in the EV membrane is co-regulated with vesicle size at the stage of biogenesis, such that the surface density of the CD63 marker remains constant.

Next, we used the Nano-SMF technique to phenotype not only CD63-positive EVs but also to probe the presence of two other tetraspanins: CD81 and CD9. EVs positive for CD9 could not be detected, which is consistent with previously reported low levels of secretion of this biomarker from HEK cells^24^ possibly in combination with poor antibody binding efficiency. CD81-positive EVs were, on the other hand, readily detected (Fig. 4i). The size distributions of the different sub-populations detected in the sample are shown in Fig. 4j-l, Fig. S17, and Table S3. We observed very low degrees of colocalization of CD63-mNeon positive and CD81 positive EVs (4%), suggesting they are discrete populations. Furthermore, the particles on which CD63 and CD81 do colocalize have a larger mean diameter (112 ± 41 nm, Fig. 4k), which indicates that they could be the result of fusion events taking place during EV release or subsequent isolation.

We find that the anti-CD81 positive EVs are in general smaller in size (70 ± 33 nm; corresponding to a size without antibody of ∼59 nm) than the wt anti-CD63 positive EVs, which may be relevant for their biofunctionality.^60^ Interestingly, the correlation between size and anti-CD81 intensity (Fig. 4l) is distinctly different than the size vs anti-CD63 or CD63-mNeon intensity correlations shown in Fig. 4h and Fig. 4b. The data suggests that CD81 content is largely independent of EV size, which contrasts the observed behavior of CD63 and therefore suggests different modus of CD81 tetraspanin sorting into EVs.

Lastly, EV analysis is highly dependent on, and often biased by, the methods that are used to isolate and concentrate these particles. An important advantage of Nano-SMF is that it requires very small sample volumes (2-20µL), and we therefore investigated the potential of detecting EVs taken directly from the cell supernatant after 48h. For this we used HEK293T cells overexpressing CD63-mCherry (Fig. 4d, Fig. S18). As reported in Table S3, we were able to estimate a concentration of 2.8•10^9^ particle/mL. This is in good agreement with the literature, considering the limited data available at this resolution.^61^ We also determined their size to 68 ± 39 nm, which is ∼ 20 nm smaller than reported in Fig. 4a. This underscores the possibility that isolation and purification steps may compromise EV integrity or introduce bias e.g. in the form of fusion events, underlining the value of analyzing EVs directly in cell culture media to better reflect their native properties.

## CONCLUSIONS

We present a nanochannel-based method that we refer to as Nano-SMF for simultaneous size determination and multiplexed fluorescence-based phenotyping of single biological nanoparticles in solution. We demonstrate that it is possible to accurately determine nanoparticle size in the 20-200 nm range based on one-dimensional Brownian motion of particles under confinement in nanochannels, captured by single particle tracking. The nanochannels are crucial for the resolution, as they keep each particle on a single track and in focus during fluorescence imaging. This greatly facilitates the multiplexed intensity analysis by allowing sequential switching of the excitation light source, but the nanochannels also provide an advantage relative to other phenotyping methods in that analyzed sample can be easily retrieved from the microreservoir at the nanochannel outlet, enabling further experimentation. The precision in size determination of Nano-SMF is on par with TEM and thus does not require the use of calibration standards as is the case in flow cytometry instrument where size is inferred from scattering intensity.^23^ Moreover, due to the spatial separation of the particles Nano-SMF has very good capacity to discriminate between small and large particles in a mixed sample, avoiding contrast problems that are common in NTA.

A significant challenge in biological nanoparticle characterization is the ability to simultaneously determine size and multiple biomarkers (via fluorescence). In recent years other nanofluidic approaches have been developed, including CliC-based methods, that employ nanoscale confinement and multiplexed fluorescence detection,^30^ increasing the diversity of available techniques and highlighting the importance of expanding the experimental toolbox for probing particles in solution. Here, we demonstrate that Nano-SMF achieves sizing and simultaneous multiplexed fluorescence detection with true single-particle discrimination, which opens new possibilities for precise characterization of the composition of individual particles in complex mixtures. We demonstrate the power of this approach by analyzing HEK cell secreted EVs and show, through antibody-based phenotyping, that the common EV biomarker tetraspanin proteins CD63 and CD81^62^ reside on distinctive sub-populations which differ in mean size. Interestingly, the abundance of CD81 in CD81-positive EVs appears largely independent on their size, whereas the abundance of CD63 in EVs from both wt and CD63-mNeon overexpressing cells scales with diameter. This reveals an intriguing insight into how these tetraspanins may be differently sorted into EVs during biogenesis and demonstrates that Nano-SMF is powerful in capturing fine-tuned, precise information on how EVs are composed under various biological conditions. Finally, the preliminary results on non-purified EVs open new perspectives for Nano-SMF to relate the impact of different factors with a minimum impact on the sample heterogeneity. This, together with the demonstrated possibility to count and characterize EVs in nanoliter volumes of conditions cell media, opens entirely new possibilities to advance EV analysis for research as well as diagnostic applications, but also opens new avenues to analyze size and compositional variations in other types of soft biological nanoparticle systems, potentially aiding research also in related fields such as drug delivery or virology.

## METHODS

### Materials

All materials were purchased from commercial sources, unless stated otherwise. 1-palmitoyl-2-oleoyl-glycero-3-phosphocholine (POPC) and fluorescent head-group labelled 1,2-dioleoylsn-glycero-3-phosphoethanolamine-N-(lissamine rhodamine B sulfonyl) (RhoB) were purchased from Avanti Polar Lipids. Fluorescent head-group labelled ATTO488 and ATTO647 1,2-Dipalmitoyl-sn-glycero-3-phosphoethanolamine (DOPE) lipids were purchased from ATTO-TEC GmbH. Marina blue 1,2-Dihexadecanoyl-*sn*-Glycero-3-Phosphoethanolamine (Marina Blue DHPE) and fluorescent polystyrene beads (Fluoro-Max red; 100 and 50 nm in diameter) were purchased from ThermoFisher Scientific. Gold (Au) nanoparticles (60 and 100 nm in diameter; OD 1; stabilized suspension in citrate buffer) were purchased from Sigma Aldrich and P-dots (StarBright Violet 670) were purchased from Bio-rad. Allophycocyanin (APC) conjugated antihuman anti-CD63 and anti-CD81 IgG primary antibodies were purchased from Miltenyi Biotec. Phosphate-buffered saline (PBS, tablets) and PKH26 Red fluorescent Cell Linker Mini Kit were purchased from Sigma-Aldrich. Illustra MicroSpin S-200 HR columns were purchased from GE Healthcare. PBS buffers were filtered with Millipore filters (0.1 mm) from Merck and water was deionized and filtered using a Milli-Q system from Merck.

### Single Particle Nanofluidic measurements

The surfaces in the nanofluidic devices were passivated against non-specific particle adsorption by formation of a POPC supported lipid bilayer (SLB) through surface-induced vesicle rupture^63^, followed by a rinsing step with PBS buffer. The SLB liposomes were doped with 1% Marina blue to allow visual control of the SLB formation by fluorescence microscopy.

Upon operating the device, sample (2-10 µL volume, with nominal concentrations of 10^8^-10^10^ particles/mL) was added to inlet 1 (Fig. 1a). A pressure of 0.5 bar was applied to inlet 1 for 10 minutes to load the sample, whereafter equal pressures of 10 mbar were applied to inlets 1 and 2 (resulting in a flow speed of 15 μm/s) to drive the sample through the nanochannels. A pressure of 30 mbar (45 μm/s) was used for Fig. 4.

Prior to use, 10 vol% SDS was added to the polystyrene bead suspension to suppress unspecific binding, in a non-passivated chip. Au nanoparticles were also examined in non-passivated channels to avoid disturbing the nanoparticles citrate stabilization; the inherent repulsive charge of the nanoparticles towards the nanochannel walls was sufficient to avoid excessive non-specific binding. Single nanoparticles in the nanochannels were tracked using epifluorescence microscopy. For all experiments, an inverted Zeiss Axio Observer Z1 microscope equipped with a 63x oil immersion objective (NA = 1.46), a Colibri 7 LED light source (Zeiss) and a Andor iXon Ultra 888 EMCCD camera (Andor Technology). FITC, TRITC, Cy5 and DAPI/FITC/TRITC/Cy5 filter cubes (CHROMA) were used.

The data was acquired with the microscope software Zen at an exposure time of 30 ms, resulting in framerates of 28.6 or 31.2 (depending on experimental setup) for single-color, 8.5 per color for dual-color and 6.8 per color for tri-color acquisition, and saved as tag image file format (.tif) videos.

### Single particle tracking analysis

The recorded microscopy videos (i.e., image time series) were analyzed using in-house MATLAB (MathWorks) scripts. Before analysis, minor image processing was done in case of (i) low signal, (ii) excessive non-specific binding to the channel surfaces and/or (iii) uneven background signal. When very low signal-to-noise ratio was observed (i), the microscopy videos were treated with a frame-wise application of a Gaussian blur filter (σ = 1 pixel) to improve the signal-to-noise ratio. In the case of excessive non-specific binding (ii), the microscopy videos were treated by subtraction of each subsequent frame. To compensate for uneven background (iii) the average of all frames was subtracted from each frame of the microscopy video.

Subsequently, the videos were analyzed with a detection algorithm^64^ that identified particle intensities and positions and linked them into tracks using a nearest neighbor linking scheme.^65^ Diffusion coefficients were determined from the mean square displacement of the particle tracks, corrected for motion blur. Hydrodynamic coupling to the nanochannel walls was accounted for by multiplication with a hinderance factor based on the channel geometry.^39,66^ Colocalization was evaluated from the overlap between particle tracks for videos acquired in multi-color and corrected for cross excitation. Concentration was determined by comparing the number of analyzed particles with the volume of the suspension passing through the field of view during acquisition. For more detailed descriptions, see Supplementary Note 1-4.

All nanoparticle sizes are given by the mean diameter ± standard deviation, both calculated from the distribution fits.

### Transmission electron microscopy (TEM)

TEM was used as reference for size determination of the 100 nm and 50 nm FluoroMax Red polystyrene beads and Au nanoparticles. Samples were drop-casted onto separate Holey Carbon Film on top of a copper grid, prod 609 (Ted Pella Inc.). After a few minutes of equilibration, excess liquid was removed, and the samples were left to dry in air and thereafter kept in a dry box until the measurement. TEM images were collected using a FEI Tecnai T20 operated at the accelerating voltage 200 kV. The size of the beads was determined using the identify circle algorithm in MATLAB (MathWorks), whereas the size of the Au nanoparticles was determined using the projected area of the particles. Aggregated particles were separated with the watershed algorithm.

### Nanoparticle Tracking Analysis (NTA)

NTA were performed using a Malvern NanoSight LM10 instrument equipped with a 488 nm laser and operating in scattering or fluorescence mode and under a flow rate of 100 (10 μL/min) obtained with a Nanosight syringe pump module and with the cameral level set to 15. Each sample was analyzed in a set of three videos of 90 seconds each. The videos were analyzed with the built-in NTA 3.2 software using a detection threshold of 3 to be able to determine optimized size distributions and concentrations. The buffer viscosity was considered as that of water at 21°C. All comparative measurements were done on the same day with the same protocol.

### Lipid vesicle preparation

Large unilamellar vesicles (LUVs) were prepared by mixing lipids at the following ratios: POPC (100 mol%), POPC/ATTO488 (98/2 mol%), POPC/Rhod (98/2 mol%), POPC/ATTO647 (98/2 mol%), POPC/ATTO488/ATTO647 (96/2/2 mol%), in round bottom flasks followed by drying of the lipid films overnight under vacuum. The lipid films were re-hydrated with PBS buffer to a total lipid concentration of 1 mg/mL. LUVs were obtained by extrusion using a mini-extruder (Avanti Polar Lipids), with two polycarbonate filters of 100 nm pore diameter (21 passages), followed by a second extrusion through filters of 30 nm pore diameter (5 passages). POPC LUVs were fluorescently labelled by mixing 10 μL of the LUV suspension with 3 μL of PKH26 dye (0.5 mM in ethanol) and 100 μL of diluent C diluting agent (provided in the labelling kit) and incubated for 10 min. Excess dye was removed using Illustra Microspin S-200 HR columns after exchanging the buffer solution of the spin columns to PBS.

### Cell culture

293 Freestyle suspension cells (ThermoFisher Scientific, FreeStyle 293-F Cells) were cultured in FreeStyle 293 Expression Medium according to manufacturer’s instructions. HEK293T cells were grown in a 1:1 mixture of minimal essential medium (MEM) and nutrient mixture F-12 Ham supplemented with 10% heat-inactivated fetal bovine serum and 1% MEM non-essential amino acids. The cells were detached (trypsin-EDTA 0.05%, 5 minutes) and passaged twice a week.

### Generation of Stable Cell lines

Codon-optimized DNA sequences coding for human CD63 (Uniprot accession number P08962) and the fluorescent proteins mNeonGreen^67^ and mCherry were synthesized (Integrated DNA Technologies) as gene fragments and cloned downstream of the CAG promoter into the pLEX vector backbone using EcoRI and NotI. To generate the different constructs expressing respective fluorescent proteins fused to the C-terminus of CD63, fluorescent protein coding sequences (CDS) were subcloned into pLEX-CD63 using SacI and NotI. Next, the complete CDS of the different CD63-fluorescent protein fusions were cloned into the lentiviral p2CL9IPwo5 backbone downstream of the SFFV promoter using EcoRI and NotI, and upstream of an internal ribosomal entry site-puromycin resistance cDNA cassette.^53^ The expression cassette was confirmed by sequencing. Respective cell lines were co-transfected with p2CL9IPw5 plasmids containing CD63 fused to the respective fluorescent proteins, the helper plasmid pCD/NL-BH, and the human codon-optimized foamy virus envelope plasmid pcoPE^68–70^ using the transfection reagent JetPEI (Polyplus, Illkrich Cedex). 16 h post transfection gene expression from the human CMV immediate-early gene enhancer/promoter was induced with 10 mM sodium butyrate (Sigma-Aldrich) for 6 h before fresh media was added to the cells, and the supernatant was collected 22 h later. Viral particles were pelleted at 25,000 × *g* for 90 min at 4°C. The supernatant was discarded, and the pellet was resuspended in 2 mL of Iscove’s Modified Dulbecco’s Media supplemented with 20% FBS and 1% P/S. Aliquots were stored at −80°C until usage. To generate stable cell lines, respective cells were transduced by overnight exposure to virus stocks and passaged at least five times under puromycin selection (Sigma; 6 µg/mL). The expression of respective CD63-fluorescent protein fusion constructs was confirmed *via* flow cytometry and fluorescence microscopy for all established cell lines (data not shown).

### Preparation and labelling of extracellular vesicles

Extracellular vesicles from the conditioned media of 293-F cells were harvested after a 48h culture period in OptiMEM (Invitrogen) as previously described.^71^ The samples were directly subjected to a low-speed centrifugation step at 500 × g for 5 min followed by a 2,000 × g spin for 10 min to remove larger particles and cell debris. Next, samples were filtered through bottle top filters (Corning, low protein binding) with cellulose acetate membranes (0.22 μm pore size) to remove any larger particles. EVs were concentrated by tangential flow filtration (TFF) with the conditioned media being diafiltrated with at least two times of the initial volume of PBS and concentrated to 20 mL using the KR2i TFF system (SpectrumLabs) equipped with modified polyethersulfone hollow fiber filters with 300 kDa membrane pore size (MidiKros, 370 cm^2^ surface area, SpectrumLabs) at a flow rate of 100 mL/min (transmembrane pressure at 3.0 psi and shear rate at 3,700 s^−1^). Finally, the EVs were concentrated to a volume of 500 μL using Amicon Ultra-15 10 kDa molecular weight cut-off spin-filters (Millipore). The particle concentration was determined to 10^12^ p/mL using NTA as described above, then kept in stabilizing PBS-HAT buffer^72^ in the freezer at -80°C aliquoted.

EVs from the conditioned media of HEK293T cells were harvested after a 48h culture period in complex media and injected (20µL) inside the chip. No further purification steps were performed.

For EV phenotyping using Nano-SMF, samples were immunolabelled under saturated staining conditions by mixing EVs (final concentration 1 × 10¹⁰ particles/mL) with antibodies (final concentration 8 nM) to a total final volume of 25 µL. The samples were thereafter incubated over night at room temperature protected from light. No additional steps were taken to separate unbound antibody from the EVs prior to analysis.

### Confocal imaging of HEK293T cells expressing CD63-mCherry

In a glass-bottom dish (Cellvis #D35C4-20-1.5-N), 30 000 cells in 500 µL of complete DMEM GlutaMAX media, (Gibco, 31966047) supplemented with 10 % FBS (Gibco), was seeded in each well and incubated for 40 h. Prior to imaging, cells were incubated with 5 µg/mL Hoechst 33342 for 30 min, washed once with serum-free medium, and imaged immediately. Imaging was performed on a Nikon Ti2 inverted microscope equipped with an X-Light V3 spinning-disk confocal unit (CrestOptics) and a Teledyne Photometrics Kinetix sCMOS camera, using a 60×/1.4 NA oil-immersion objective. Hoechst 33342 and CD63–mCherry were excited at 405 nm and 546 nm, respectively (Lumencor Celesta), and emission was collected using bandpass filters at 438/24 nm (Hoechst) and 595/31 nm (mCherry). All imaging was conducted using a stage-top incubator (UNO-T system, Okolab, Italy) maintaining cells at 37 °C and 5% CO₂.

## ACKNOWLEDGEMENTS

This project has been funded by grants from the Knut and Alice Wallenberg Foundation (KAW projects 2015.0055 and 2022.0134, and Wallenberg Academy Fellow program 2019.0238, the Swedish Foundation for Strategic Research (SSF) via the FoRmulaEx industrial research centre for RNA delivery (IRC15-0065), and the Wenner-Gren foundation postdoc program. The nanofluidic devices used in this work were fabricated at Chalmers MyFab cleanroom facility.

## Notes

### Competing Interest Statement

The authors have declared no competing interest.

## REFERENCES

1. Van Niel, G., D’Angelo, G. & Raposo, G. Shedding light on the cell biology of extracellular vesicles. Nat. Rev. Mol. Cell Biol. 19, 213–228 (2018).

2. Colombo, M., Raposo, G. & Théry, C. Biogenesis, Secretion, and Intercellular Interactions of Exosomes and Other Extracellular Vesicles. Annu. Rev. Cell Dev. Biol. 30, 255–289 (2014).

3. Van Niel, G. et al. Challenges and directions in studying cell–cell communication by extracellular vesicles. Nat. Rev. Mol. Cell Biol. 23, 369–382 (2022).

4. Möller, A. & Lobb, R. J. The evolving translational potential of small extracellular vesicles in cancer. Nat. Rev. Cancer 20, 697–709 (2020).

5. Yuana, Y., Sturk, A. & Nieuwland, R. Extracellular vesicles in physiological and pathological conditions. Blood Rev. 27, 31–39 (2013).

6. Murphy, D. E. et al. Extracellular vesicle-based therapeutics: natural versus engineered targeting and trafficking. Exp. Mol. Med. 51, (2019).

7. El Andaloussi, S., Mäger, I., Breakefield, X. O. & Wood, M. J. A. Extracellular vesicles: biology and emerging therapeutic opportunities. Nat. Rev. Drug Discov. 12, 347–357 (2013).

8. El-Andaloussi, S. et al. Exosome-mediated delivery of siRNA in vitro and in vivo. Nat. Protoc. 7, 2112–2126 (2012).

9. Liang, X., et al. Engineering of extracellular vesicles for efficient intracellular delivery of multimodal therapeutics including genome editors. Preprint at 10.21203/rs.3.rs-3329019/v1 (2023).

10. Zheng, W. et al. Identification of scaffold proteins for improved endogenous engineering of extracellular vesicles. Nat. Commun. 14, (2023).

11. Nordin, J. Z. et al. Ultrafiltration with size-exclusion liquid chromatography for high yield isolation of extracellular vesicles preserving intact biophysical and functional properties. *Nanomedicine Nanotechnol*. Biol. Med. 11, 879–883 (2015).

12. Smith, Z. J., et al. Single exosome study reveals subpopulations distributed among cell lines with variability related to membrane content. J. Extracell. Vesicles 4, (2015).

13. Welsh, J. A., et al. Minimal information for studies of extracellular vesicles (MISEV2023): From basic to advanced approaches. J. Extracell. Vesicles 13, e12404 (2024).

14. Willms, E., Cabañas, C., Mäger, I., Wood, M. J. A. & Vader, P. Extracellular Vesicle Heterogeneity: Subpopulations, Isolation Techniques, and Diverse Functions in Cancer Progression. Front. Immunol. 9, (2018).

15. Sork, H. et al. Heterogeneity and interplay of the extracellular vesicle small RNA transcriptome and proteome. Sci. Rep. 8, (2018).

16. Willms, E. et al. Cells release subpopulations of exosomes with distinct molecular and biological properties. Sci. Rep. 6, (2016).

17. Gurunathan, S., Kang, M.-H., Jeyaraj, M., Qasim, M. & Kim, J.-H. Review of the Isolation, Characterization, Biological Function, and Multifarious Therapeutic Approaches of Exosomes. Cells 8, 307 (2019).

18. Kowal, J. et al. Proteomic comparison defines novel markers to characterize heterogeneous populations of extracellular vesicle subtypes. Proc. Natl. Acad. Sci. 113, (2016).

19. Zabeo, D., et al. Exosomes purified from a single cell type have diverse morphology. J. Extracell. Vesicles 6, (2017).

20. Dragovic, R. A. et al. Sizing and phenotyping of cellular vesicles using Nanoparticle Tracking Analysis. Nanomedicine Nanotechnol. Biol. Med. 7, 780–788 (2011).

21. Dragovic, R. A. et al. Isolation of syncytiotrophoblast microvesicles and exosomes and their characterisation by multicolour flow cytometry and fluorescence Nanoparticle Tracking Analysis. Methods 87, 64–74 (2015).

22. Görgens, A., et al. Optimisation of imaging flow cytometry for the analysis of single extracellular vesicles by using fluorescence-tagged vesicles as biological reference material. J. Extracell. Vesicles 8, (2019).

23. Tian, Y. et al. Protein Profiling and Sizing of Extracellular Vesicles from Colorectal Cancer Patients *via* Flow Cytometry. ACS Nano 12, 671–680 (2018).

24. Daaboul, G. G. et al. Digital Detection of Exosomes by Interferometric Imaging. Sci. Rep. 6, (2016).

25. Mitra, A., Deutsch, B., Ignatovich, F., Dykes, C. & Novotny, L. Nano-optofluidic Detection of Single Viruses and Nanoparticles. ACS Nano 4, 1305–1312 (2010).

26. Penders, J. et al. Single Particle Automated Raman Trapping Analysis of Breast Cancer Cell-Derived Extracellular Vesicles as Cancer Biomarkers. ACS Nano 15, 18192–18205 (2021).

27. Cho, S. et al. Multifluorescence Single Extracellular Vesicle Analysis by Time-Sequential Illumination and Tracking. ACS Nano 15, 11753–11761 (2021).

28. Penders, J. et al. Single Particle Automated Raman Trapping Analysis. Nat. Commun. 9, (2018).

29. Kashkanova, A. D., Blessing, M., Gemeinhardt, A., Soulat, D. & Sandoghdar, V. Precision size and refractive index analysis of weakly scattering nanoparticles in polydispersions. Nat. Methods 19, 586–593 (2022).

30. Boateng, E. et al. Multiparametric Characterization of Individual Suspended Nanoparticles Using Confocal Fluorescence and Interferometric Scattering Microscopy with Microfluidic Confinement. Nano Lett. 25, 12158–12165 (2025).

31. Kamanzi, A. et al. Single-Particle Multiparametric Microscopy Reveals Structural, Size, and Payload Heterogeneity in mRNA-Loaded Lipid Nanoparticles. ACS Nano 20, 1290–1303 (2026).

32. Fujise, K., Mishra, J., Rosenfeld, M. S. & Rafiq, N. M. Synaptic vesicle characterization of iPSC-derived dopaminergic neurons provides insight into distinct secretory vesicle pools. Npj Park. Dis. 11, 16 (2025).

33. Müller, V. & Westerlund, F. Optical DNA mapping in nanofluidic devices: principles and applications. Lab. Chip 17, 579–590 (2017).

34. Frykholm, K., Nyberg, L. K. & Westerlund, F. Exploring DNA–protein interactions on the single DNA molecule level using nanofluidic tools. Integr. Biol. 9, 650–661 (2017).

35. Friedrich, R. et al. A nano flow cytometer for single lipid vesicle analysis. Lab. Chip 17, 830–841 (2017).

36. Block, S., Fast, B. J., Lundgren, A., Zhdanov, V. P. & Höök, F. Two-dimensional flow nanometry of biological nanoparticles for accurate determination of their size and emission intensity. Nat. Commun. 7, (2016).

37. Van Der Pol, E., Van Gemert, M. J. C., Sturk, A., Nieuwland, R. & Van Leeuwen, T. G. Single vs. swarm detection of microparticles and exosomes by flow cytometry. J. Thromb. Haemost. 10, 919–930 (2012).

38. Block, S., Fast, B. J., Lundgren, A., Zhdanov, V. P. & Höök, F. Two-dimensional flow nanometry of biological nanoparticles for accurate determination of their size and emission intensity. Nat. Commun. 7, (2016).

39. Dechadilok, P. & Deen, W. M. Hindrance Factors for Diffusion and Convection in Pores. Ind. Eng. Chem. Res. 45, 6953–6959 (2006).

40. Arab, T., et al. Characterization of extracellular vesicles and synthetic nanoparticles with four orthogonal single-particle analysis platforms. J. Extracell. Vesicles 10, (2021).

41. Faez, S. et al. Fast, Label-Free Tracking of Single Viruses and Weakly Scattering Nanoparticles in a Nanofluidic Optical Fiber. ACS Nano 9, 12349–12357 (2015).

42. Caputo, F. et al. Measuring particle size distribution and mass concentration of nanoplastics and microplastics: addressing some analytical challenges in the sub-micron size range. J. Colloid Interface Sci. 588, 401–417 (2021).

43. Gang, H. et al. Microfluidic Diffusion Platform for Characterizing the Sizes of Lipid Vesicles and the Thermodynamics of Protein–Lipid Interactions. Anal. Chem. 90, 3284–3290 (2018).

44. Fraikin, J.-L., Teesalu, T., McKenney, C. M., Ruoslahti, E. & Cleland, A. N. A high-throughput label-free nanoparticle analyser. Nat. Nanotechnol. 6, 308–313 (2011).

45. Matsuura, Y., Nakamura, A. & Kato, H. Determination of Nanoparticle Size Using a Flow Particle-Tracking Method. Anal. Chem. 90, 4182–4187 (2018).

46. Špačková, B. et al. Label-free nanofluidic scattering microscopy of size and mass of single diffusing molecules and nanoparticles. Nat. Methods 19, 751–758 (2022).

47. Filipe, V., Hawe, A. & Jiskoot, W. Critical Evaluation of Nanoparticle Tracking Analysis (NTA) by NanoSight for the Measurement of Nanoparticles and Protein Aggregates. Pharm. Res. 27, 796–810 (2010).

48. Jõemetsa, S. et al. Independent Size and Fluorescence Emission Determination of Individual Biological Nanoparticles Reveals that Lipophilic Dye Incorporation Does Not Scale with Particle Size. Langmuir 36, 9693–9700 (2020).

49. Nele, V. et al. Effect of Formulation Method, Lipid Composition, and PEGylation on Vesicle Lamellarity: A Small-Angle Neutron Scattering Study. Langmuir 35, 6064–6074 (2019).

50. Van Der Vlist, E. J., Nolte-’t Hoen, E. N. M., Stoorvogel, W., Arkesteijn, G. J. A. & Wauben, M. H. M. Fluorescent labeling of nano-sized vesicles released by cells and subsequent quantitative and qualitative analysis by high-resolution flow cytometry. Nat. Protoc. 7, 1311–1326 (2012).

51. Lubart, Q. et al. Lipid vesicle composition influences the incorporation and fluorescence properties of the lipophilic sulphonated carbocyanine dye SP-DiO. Phys. Chem. Chem. Phys. 22, 8781–8790 (2020).

52. Dehghani, M., Gulvin, S. M., Flax, J. & Gaborski, T. R. Systematic Evaluation of PKH Labelling on Extracellular Vesicle Size by Nanoparticle Tracking Analysis. Sci. Rep. 10, 9533 (2020).

53. Wiklander, O. P. B. et al. Systematic Methodological Evaluation of a Multiplex Bead-Based Flow Cytometry Assay for Detection of Extracellular Vesicle Surface Signatures. Front. Immunol. 9, (2018).

54. Hurwitz, S. N., Conlon, M. M., Rider, M. A., Brownstein, N. C. & Meckes, D. G. Nanoparticle analysis sheds budding insights into genetic drivers of extracellular vesicle biogenesis. J. Extracell. Vesicles 5, (2016).

55. Hurwitz, S. N. et al. CD63 Regulates Epstein-Barr Virus LMP1 Exosomal Packaging, Enhancement of Vesicle Production, and Noncanonical NF-κB Signaling. J. Virol. 91, (2017).

56. Paulaitis, M., Agarwal, K. & Nana-Sinkam, P. Dynamic Scaling of Exosome Sizes. Langmuir 34, 9387–9393 (2018).

57. Tan, Y. H. et al. A Nanoengineering Approach for Investigation and Regulation of Protein Immobilization. ACS Nano 2, 2374–2384 (2008).

58. Brennan, K. et al. A comparison of methods for the isolation and separation of extracellular vesicles from protein and lipid particles in human serum. Sci. Rep. 10, (2020).

59. Sung, B. H. et al. A live cell reporter of exosome secretion and uptake reveals pathfinding behavior of migrating cells. Nat. Commun. 11, (2020).

60. Lee, S.-S. et al. A novel population of extracellular vesicles smaller than exosomes promotes cell proliferation. Cell Commun. Signal. 17, (2019).

61. Lerner, N., Avissar, S. & Beit-Yannai, E. Extracellular vesicles mediate signaling between the aqueous humor producing and draining cells in the ocular system. PLOS ONE 12, e0171153 (2017).

62. Jeppesen, D. K. et al. Reassessment of Exosome Composition. Cell 177, 428–445.e18 (2019).

63. Persson, F. et al. Lipid-Based Passivation in Nanofluidics. Nano Lett. 12, 2260–2265 (2012).

64. Block, S., Glöckl, G., Weitschies, W. & Helm, C. A. Direct Visualization and Identification of Biofunctionalized Nanoparticles using a Magnetic Atomic Force Microscope. Nano Lett. 11, 3587–3592 (2011).

65. Conn, P. M. Imaging and Spectroscopic Analysis of Living Cells. (Academic Press, San Diego, Calif, 2012).

66. Van Der Sman, R. G. M. Drag force on spheres confined on the center line of rectangular microchannels. J. Colloid Interface Sci. 351, 43–49 (2010).

67. Shaner, N. C. et al. A bright monomeric green fluorescent protein derived from Branchiostoma lanceolatum. Nat. Methods 10, 407–409 (2013).

68. Mochizuki, H., Schwartz, J. P., Tanaka, K., Brady, R. O. & Reiser, J. High-Titer Human Immunodeficiency Virus Type 1-Based Vector Systems for Gene Delivery into Nondividing Cells. J. Virol. 72, 8873–8883 (1998).

69. Leurs, C. et al. Comparison of Three Retroviral Vector Systems for Transduction of Nonobese Diabetic/Severe Combined Immunodeficiency Mice Repopulating Human CD34^+^Cord Blood Cells. Hum. Gene Ther. 14, 509–519 (2003).

70. Müllers, E. et al. Novel Functions of Prototype Foamy Virus Gag Glycine- Arginine-Rich Boxes in Reverse Transcription and Particle Morphogenesis. J. Virol. 85, 1452–1463 (2011).

71. Corso, G. et al. Reproducible and scalable purification of extracellular vesicles using combined bind-elute and size exclusion chromatography. Sci. Rep. 7, (2017).

72. Görgens, A., et al. Identification of storage conditions stabilizing extracellular vesicles preparations. J. Extracell. Vesicles 11, e12238 (2022).

