## Supporting Information for "SIZE DETERMINATION AND MULTIPLEXED FLUORESCENCE-BASED PHENOTYPING OF SINGLE CELL-DERIVED MEMBRANE VESICLES USING A NANOFLUIDIC DEVICE"

Current affiliations:

[†]Sciflux OÜ, Harku, Estonia

[‡]Michael Smith Laboratories, University of British Columbia, Vancouver, BC V6T 1Z4, Canada

*Corresponding authors:

Elin K. Esbjörner,

Fredrik Westerlund,

### 1. Supplementary Notes

#### Supplementary Note 1: Single particle tracking (SPT)

The collected microscopy videos were analyzed with in-house MATLAB scripts. The position of potential particles (e.g., beads, liposomes and extracellular vesicles) were identified in the videos based on a detection algorithm[1], which identified local intensity maxima within each frame and stored the detected position if the intensity of the local maximum exceeded a user defined threshold (which was determined by visual inspection of the video in ImageJ)[2]. This yielded a list of potential particles, containing information on the particle's position (frame number and pixel-resolved $x$- and $y$-coordinate) as well as the intensity of its brightest pixel. Most entries in this list are generated by real particles, while a small fraction is created by false-positive detection due to pixel noise that exceeded by chance the intensity threshold used in the detection process. These falsely detected particles are removed in a later stage of the data analysis by application of a 2D Gaussian fit to the intensity profile of a potential particle (discussed in further detail below).

The motion of the detected particles was tracked across the frames based on a nearest neighbor linking scheme[3]: In order to identify the motion of a particular particle between 2 successive frames, the distance of the initial particle position (starting frame) to the positions of all particles detected in the successive frame was calculated, allowing to identify the particle position in the successive frame that displayed the smallest distance from the initial position. These two positions were then linked, yielding trajectories of the particle's motion throughout the video. This process required to define a distance cutoff value, above which a link of two particle is regarded to be unreasonably large (i.e., at which the motion of the particle was larger than physically possible). The validity of the distance cutoff value used was verified by a visual comparison of the extracted tracks and the original video.

After extraction of the particle trajectories, sub-pixel positions of all tracked particles were localized by application of a 2D Gaussian fit to the measured intensity profiles. The starting values of this fitting procedure were given by the (pixel-resolved) $x$- and $y$-coordinate of the to-be-fitted particle, while the width of the 2D Gaussian was fixed to 3 pixels, which roughly corresponds to the width of the point spread function of a point emitter (for the optical configuration used). The starting values of this fitting procedure were given by the (pixel-resolved) x- and $y$-coordinate of the to-be-fitted particle, while the width of the 2D Gaussian was fixed to 3 pixels, which roughly corresponds to the width of the point spread function of a point emitter (for the optical configuration used). Applied to experimental data, this fitting procedure allowed us to resolve the particles' sub-pixel position with typical localization accuracies being better than 10% of the pixel size. Applied to falsely detected particles (i.e., to pixels that were detected as their intensity noise exceeding by chance the intensity threshold), this fitting procedure typically failed, and the corresponding entry was marked and not used for further data analysis.

In the final step of the data analysis, the total intensity of the particles was extracted[4]. This was achieved on a frame-by-frame basis by cropping out, for each detected particle, a 41 pixel x 41 pixel region from the entire frame that is centered on the particle under investigation. As the particle to background ratio in our setup is low, most pixels are not associated with any particle, and the background intensity can be extracted conveniently by calculating the median intensity value of the investigated region. As the particle to background ratio in our setup is low, most pixels are not associated to any particle, and the background intensity can be extracted conveniently by calculating the median intensity value of the investigated region. This information was then used to identify all pixels being associated to the particle under investigation, which form a connected area containing all pixels around and including the detected sub-pixel coordinate, with intensity above the determined background level. The total particle intensity was determined as the sum of the intensities of these pixels and corrected for the background intensity (calculated by the product of the number of pixels times the background intensity value per pixel determined as described above).

After this analysis step, the particle tracking data was available for further analysis, which contained information on the particles' $x$- and $y$-coordinates (pixel- and subpixel-resolved values) as well as their total (integrated) fluorescence intensity. Finally, the deviation between pixel- as well as subpixel-resolved values was calculated, the distribution of which typically follows a Gaussian distribution being centered around 0 and having a standard deviation of approximately one pixel. All subpixel-resolved values that showed a larger deviation than one pixel from the corresponding pixel-resolved values were marked and rejected from further analysis, as such pronounced deviations are indicative for errors done in the localization process (e.g., breakdown of the Gaussian distribution due to too close proximity of two particles or failure in the fitting process due to too weak particle intensity). Hence, this filtering step rejects predominantly unreliable particle positions and thus improves the accuracy of the data analysis.

#### Supplementary Note 2: Size determination

The motion of the tracked particles was decomposed into stochastic (diffusion) and deterministic (drift in the flow direction) motional components as recently described[4]. In brief, if $(x_i, y_i)$ denotes the $i$-th position of a particular trajectory (covering $N$ frames in total and being separated by a time period of $\Delta t$), the drift velocity in the $x$- and $y$-directions ($v_x$ and $v_y$, respectively) was determined independently by fitting a linear function to the time-dependence of the particle's $x$- and $y$-coordinate, $(i \cdot \Delta t, x_i)$ and $(i \cdot \Delta t, y_i)$, for $i$ ranging between 1 and $N$. The stochastic contribution to the motion and $x$- and $y$-direction was extracted based on the extraction of the mean squared displacement using

$$MSD_x(n \cdot \Delta t) = var(x_i - x_{i+n}) \text{ and } MSD_y(n \cdot \Delta t) = var(y_i - y_{i+n}),$$

in which the variance $var$ was calculated using all spatial displacements $x_i - x_{i+n}$ and $y_i - y_{i+n}$ that connected particle positions being displaced by $n$ frames and that were addressable for $i$ ranging between 1 and $N - n$. For $n \ll N$, the one-dimensional diffusion coefficient in the $x$- or $y$-direction ($D_x$ and $D_y$, respectively) can be calculated using

$$D_x = MSD_x(n \cdot \Delta T)/(2 \cdot n \cdot \Delta T \cdot \xi) \text{ and } D_y = MSD_y(n \cdot \Delta T)/(2 \cdot n \cdot \Delta T \cdot \xi),$$

with $\xi$ denoting a correction factor that accounts for the motional blur effect caused by the non-zero exposure time $\Delta t_e$ of the camera[5]:

$$\xi = n - \Delta t_e/(3 \cdot \Delta t).$$

As the particles were confined in nanochannels and as the nanochannels were typically aligned with respect to the $y$-direction, no drift was detectable in the $x$-direction ($v_x \approx 0$). Furthermore, the drift velocity in the $y$-direction was, for the particle sizes used in this study, essentially indicative for the flow velocity in the middle of the nanochannels and thus did also not contain useful information for size determination of the tracked particles. Hence, the drift velocities were not used for further analysis.

In contrast, the diffusion coefficient in the $y$-direction ($D_y$; parallel to the nanochannel) depends on the particle size and can thus be used to extract size distributions. For a spherical particle moving in the bulk, the particle's diffusion coefficient $D_{bulk}$ is given by the Stokes-Einstein equation

$$D_{bulk} = k_B T/(6 \cdot \pi \cdot \eta \cdot r)$$

with $k_B T$ denoting the thermal energy (i.e., the product of Boltzmann's constant $k_B$ and the absolute termperature $T$), $\eta$ the viscosity of the bulk solution (≈ 1 mPa s for water at 20°C) and $r$ the hydrodynamic radius of the particle[6]. Particles moving in the nanochannels, however, experience an additional hydrodynamic coupling to the walls, which causes the nanoparticle's diffusion coefficient $D$ to decrease below its bulk value $D_{bulk}$ (Fig. S2)[7]. This behavior was accounted for by introducing a hindrance factor $H$ according to

$$D = H \cdot D_{bulk}.$$

For a nanochannel with a squared cross-section and the half edge length $a$, this hindrance factor has been numerically determined using the so-called center-line approximation to be

$$H = \frac{1 - 2.105 \cdot \lambda + 2.0865 \cdot \lambda^3 - 1.7068 \cdot \lambda^5 + 0.72603 \cdot \lambda^6}{1 - 0.666 \cdot \lambda^2 - 0.20217 \cdot \lambda^5}$$

using the dimensionless parameter $\lambda = \sqrt{\pi} \cdot r/a$, which depends only on the hydrodynamic radius of the particle and the nanochannel edge length[8]. Based on this relationship, the expected diffusion coefficient $D(r)$ of particles moving in the nanochannel could be calculated as a function of $r$ and as this relationship is monotonous, it could be numerically inverted to yield the function $r(D)$, allowing to extract the particle's hydrodynamic radius $r$ based on the measured diffusion coefficient in $y$-direction $D_y$. For particle radii ranging between 25 and 75 nm (which is the range investigated in this study), the bulk diffusion coefficient in water would range between 2.9 μm$^2$/s ($r$ = 75 nm) and 8.8 μm$^2$/s ($r$ =25 nm). For motion of the particles in nanochannels with an edge length of 300 nm, the hindrance factors range between 0.26 ($r$ =75 nm) and 0.71 ($r$ =25 nm), so that the diffusion coefficient in the nanochannels reduces to 0.76 μm$^2$/s ($R$ =75 nm) and 6.3 μm$^2$/s ($r$ =25 nm).

In principle, the diffusion coefficient perpendicular to the nanochannels, $D_x$, could also be used for such a size extraction. Nevertheless, even at the high acquisition rates used in this study ($\Delta t \approx$ 30 ms), the large particles ($r$ = 75 nm) will already have bounced at least once with the nanochannel walls during the acquisition of an image, so that the extracted $D_x$ is not indicative for the true diffusion coefficient of the particles and could therefore also not be used for size determination.

Finally, the size distribution was fitted with a gaussian or log-normal distribution. The median and standard deviation (σ) were used to express the size.

#### Supplementary Note 3: Colocalization and cross excitation correction

To detect colocalization between different color channels, the particle trajectories obtained from analysis of each individual color channel (two or three) were compared. Particles were considered colocalized if found within a few pixels from each other for a set number of frames (in general about 2/3 of the minimum track length used for analysis). When performing multicolor fluorescence imaging, cross excitation effects can occur resulting in emission of fluorescent dyes being detected in multiple color channels (Fig. S8). Where crosstalk was observed, particles with intensity ratio of 0.4 or less between color channels with expected crosstalk (Fig. S) were excluded in the dimmer color channel. In Fig. S9, we can detect ATTO488/ATTO647 labelled LUVs (blue/red) in the green channel, from excitation at 550 nm (Fig. S9b). To assess this cross excitation, we plotted intensity vs intensity for each set of two colors (Fig S10Supplementary Figure a). When doing so, only $I_{blue}$ plotted against $I_{red}$ followed a linear relation with an intensity ratio close to 1, which is expected for the LUVs containing equal amounts of the two dyes (ATTO488/ATTO647). In contrast, for $I_{green}$ plotted against $I_{red}$ the intensity ratio was far lower than 1 due to the fact that emission intensity in the green channel only derives from cross excitation. To separate these two populations, a linear cutoff was applied to remove all particles that were disproportionally dim in one color channel, which were considered cross excitation artefacts (Fig. S10Supplementary Figure b). By applying such a cutoff, we could obtain the particles hydrodynamic size distribution for each channel after cross excitation correction (compare Fig. S9 and Fig. S11). Fig. S12 presents the same case but in a sample with a mix of ATTO448/ATTO647 labelled LUVs (blue/red), and Rhodamine labelled LUVs (green). We detected cross excitation in the green channel from the red channel (Fig. S12e), but also from the green channel to the blue channel (Fig S12d). Again, by plotting intensity vs intensity for each set of two colors we could determine a cutoff to separate particles colocalizing from the cross excitation (Fig. S13), resulting in the graphs in Fig. S14. The particles hydrodynamic size distribution for each channel after cross excitation correction are displayed in Fig. 3c-f.

#### Supplementary Note 4: Concentration estimation

From the particle tracking, the concentration of particles was estimated through the simple formula $c = N/V$, where $c$ is concentration, $N$ is the number of particles observed and $V$ is the total volume of solution that passes through the field of view. $V = v_{flow} \cdot t \cdot w_{nc} \cdot h_{nc} \cdot n_{nc}$ is given by the mean flow speed of all tracked particles ($v_{flow}$), time of the experiment ($t$), nanochannel dimensions (width $w_{nc}$ and height $h_{nc}$) and number of nanochannels in the field of view ($n_{nc}$). To avoid the risk of counting particles twice, $N$ was determined as all particles observed in a slice with the width of the distance cutoff in the center of the field of view.

#### 2. Supplementary Data

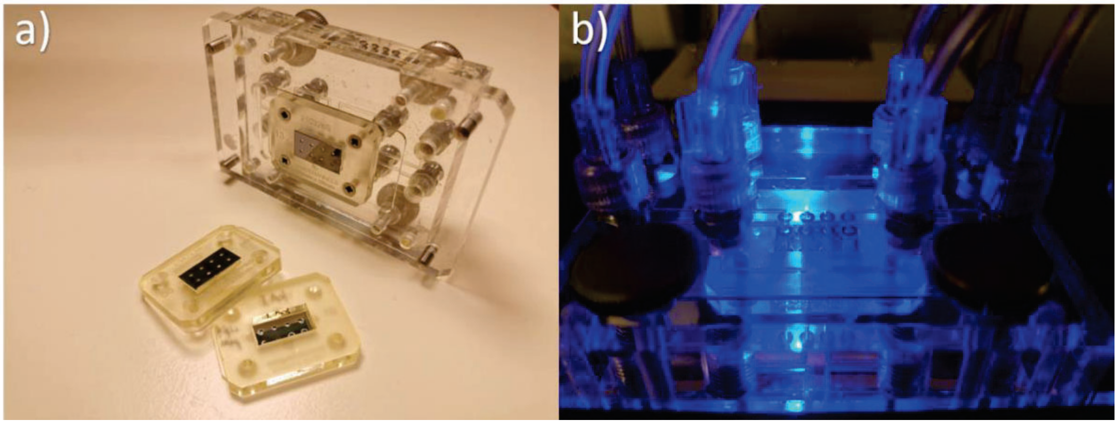

*Supplementary Figure 1: Nanofluidic chips, one attached to a holder for mounting on an epi-fluorescence microscope. a) Image of three nanofluidic chips, where one is attached to a chip holder. b) Nanofluidic chip mounted on an epi-fluorescence microscope. Attached tubing is for control of* $N_2$ *pressure for manipulating the flow inside the chip.*

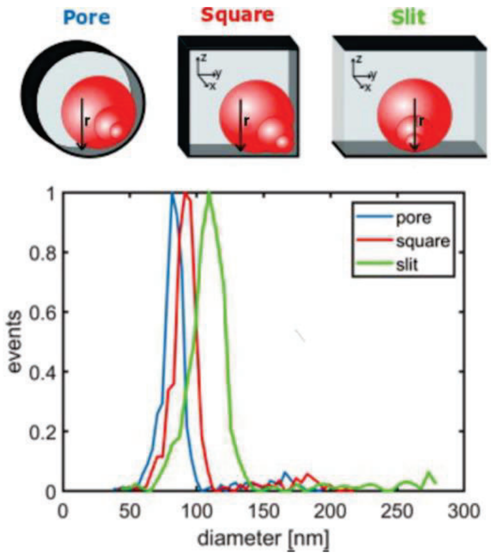

*Supplementary Figure 2. An overview of how hinderance factor depends on nanochannel geometry. Illustrations of a pore, a square and a slit shaped nanochannel in the plane perpendicular to the flow. The graph displays an example of size determination using hindrance factor equation for a pore (blue line), square (red line) and a slit (green line).*

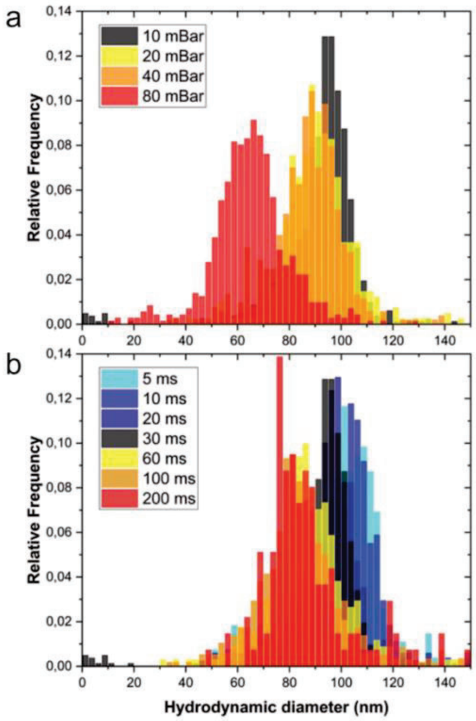

*Supplementary Figure 3. Size histograms for polystyrene fluorescent beads with (a) increasing flow pressure (i.e. flow rates corresponding to: 10 mBar ≈ 15 µm/s, 20 mBar ≈ 30 µm/s, 40 mBar ≈ 60 µm/s and 80 mBar ≈ 120 µm/s) or (b) increasing exposure times. All measurements in a) were done at an exposure time of 30 ms and all measurements in (b) were done at a flow pressure of 10 mBar.*

As mentioned earlier, both flow rate and exposure time can affect the determination of the size and if not kept sufficiently low they will impact the determined size (Fig. S3Supplementary Figure 3). Measurements were therefore done at a flow speed of 10 mbar and an exposure time of 30 ms throughout this work.

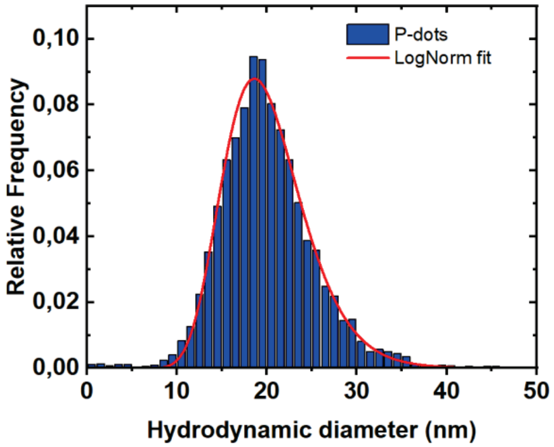

*Supplementary Figure 4. Size histogram of 20 nm P-dots characterized with Nano-SMF. Fitted with a log-normal.*

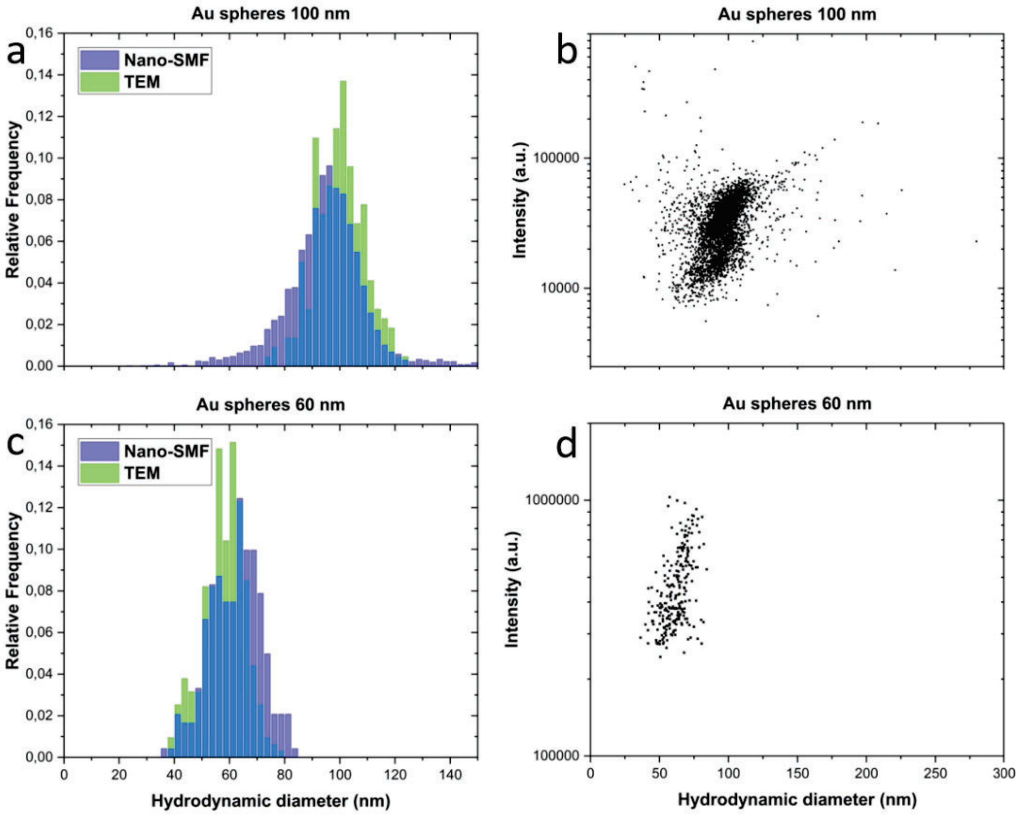

*Supplementary Figure 5. Sizes of 100 nm and 60 nm Au nanoparticles determined with Nano-SMF and TEM. a) Size histogram of 100 nm Au spheres. b) Scattering intensity vs size determined with Nano-SMF for 100 nm Au spheres. c) Size histogram of 60 nm Au spheres. d) Scattering intensity vs size determined with Nano-SMF for 60 nm Au spheres.*

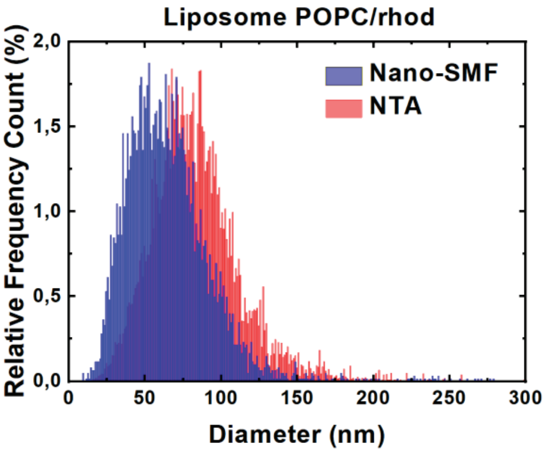

*Supplementary Figure 6. Size histogram comparison of POPC LUVs containing 2% rhodamine-labelled DHPE characterized with Nano-SMF (blue) and NTA (red).*

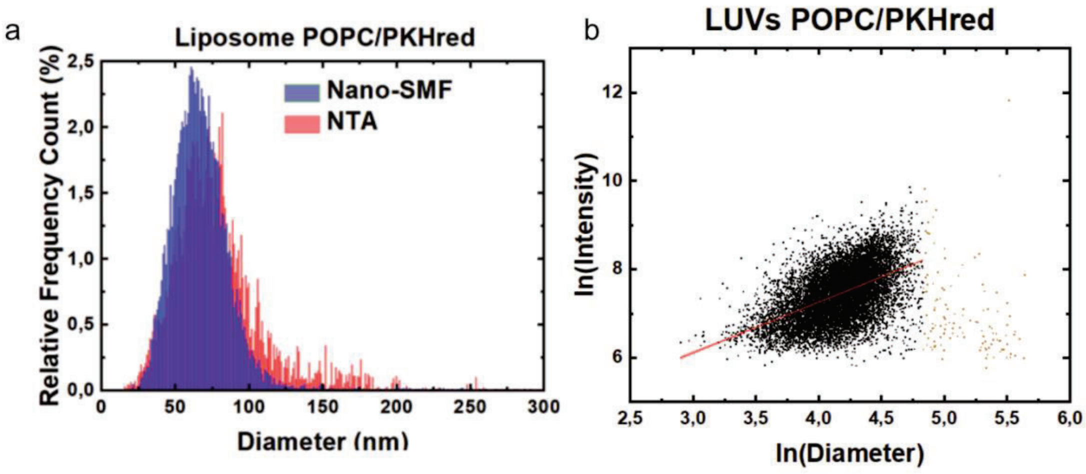

*Supplementary Figure 7. a) Size histogram comparison of POPC LUVs labelled with the lipophilic dye PKHred characterized with Nano-SMF (blue) and NTA (red). b) Logarithmic scatter plot of intensity vs diameter measured with Nano-SMF for LUVs POPC/PKHred with linear fit value X = 1.15 ± 0.02 for the bulk part of the population (black).*

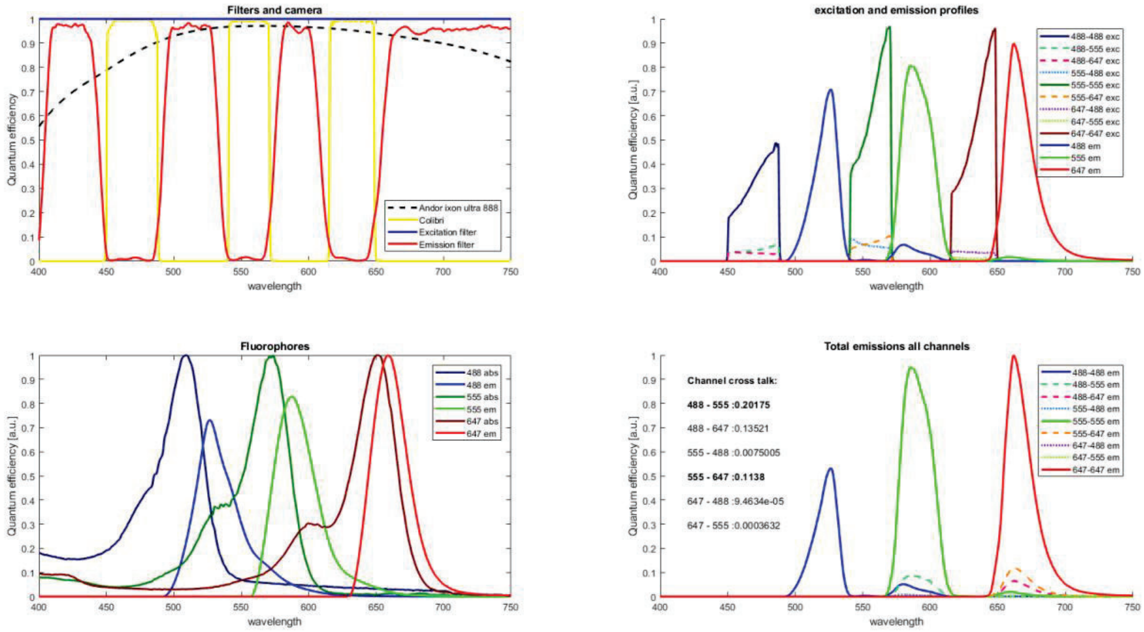

*Supplementary Figure 8. Overview of filters and excitation and emission profiles of the experimental setup and the fluorescent probes. a) Excitation and emission filters of our microscopy setup as well as excitation expressed by the Colibri light source and the quantum efficiency profile of the camera. b) Normalized absorption and emission spectra of the probes used. The height of the emission spectra corresponds to the respective quantum efficiency and amount of staining. c) Excitation and emission profiles between each color channel. d) Total expected emission seen in each color channel for excitation in each color channel.*

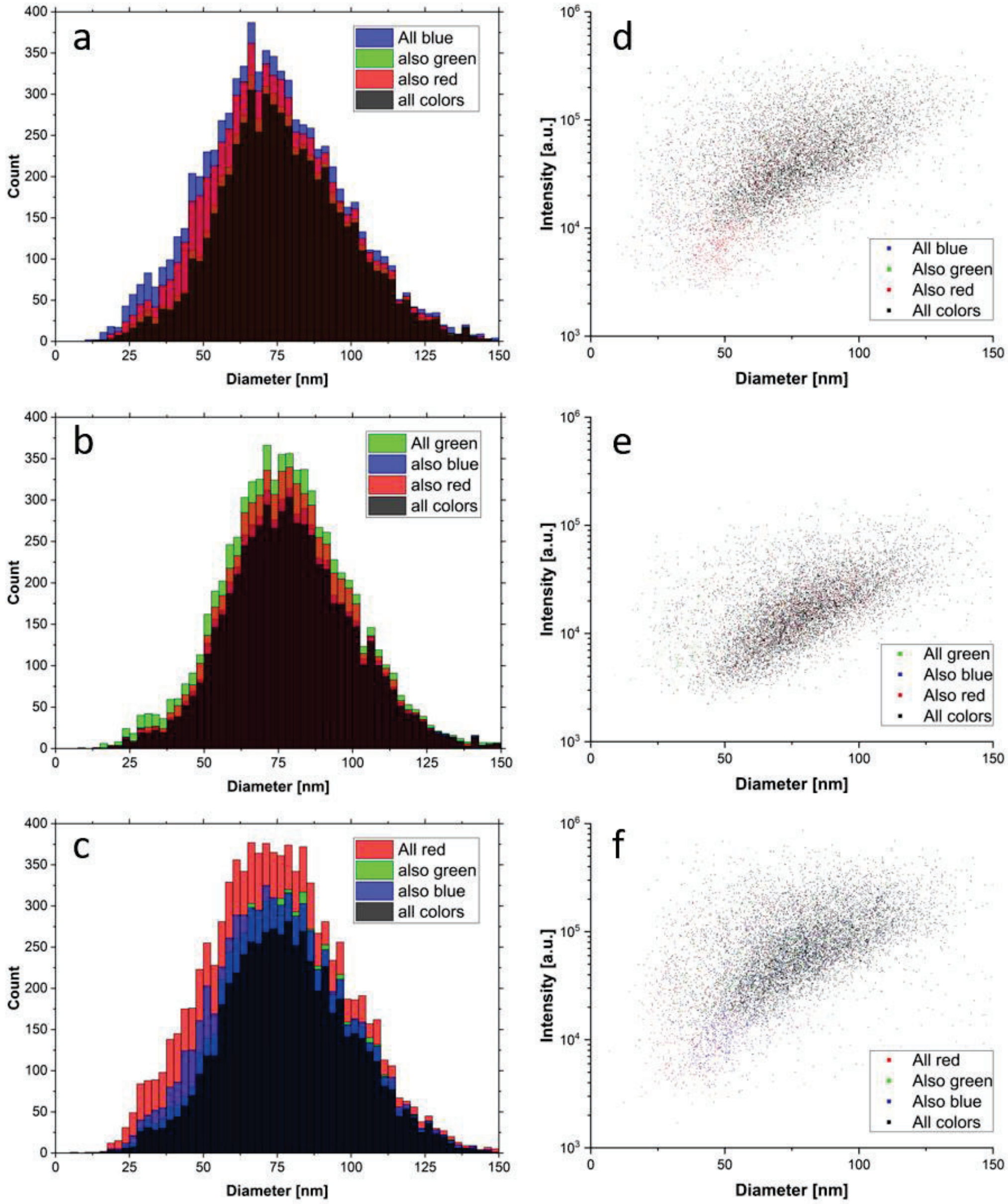

*Supplementary Figure 9. Demonstration of colocalization with Nano-SMF for a sample of LUVs POPC/(ATTO488 and ATTO647) without crosstalk correction. To the left are size histograms from the three color channels: a) blue (488 nm), b) green (550 nm) and c) red (647 nm). To the right are scatter plots of intensity vs diameter from color channel: d) blue (488 nm), e) green (550 nm) and f) red (647 nm).*

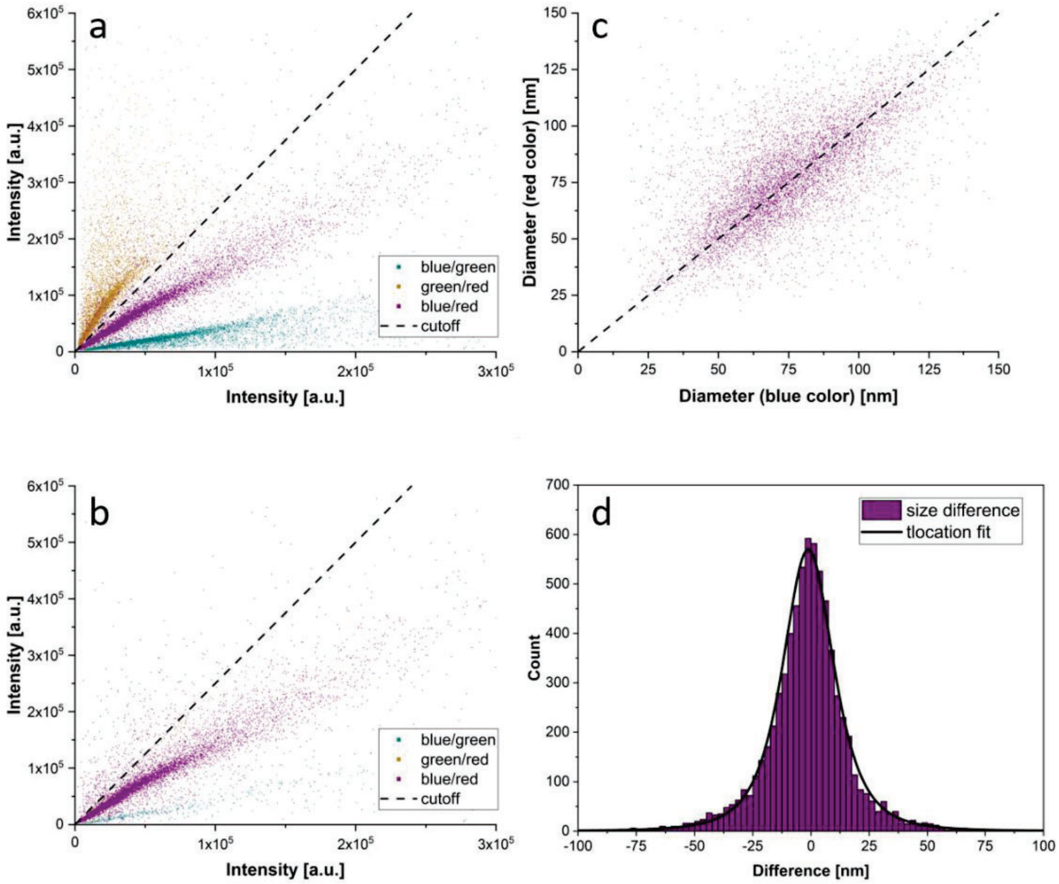

*Supplementary Figure 10. Additional characterization for the colocalization of LUVs POPC/(ATTO488 and ATTO647). a) Scatter plot of intensities from one color vs another. Crosstalk is visible in the form of a steep gradient in correlation between blue/green and green/red color channels. b) Same as (a) but corrected for crosstalk by the cutoff 0.4 (dashed black line). c) Comparison between sizes determined in blue color channel vs in red color channel. d) Histogram of size difference between sizes determined in blue and red. The t-location fit parameters $\sigma$=11.6 and $\mu$=2.8.*

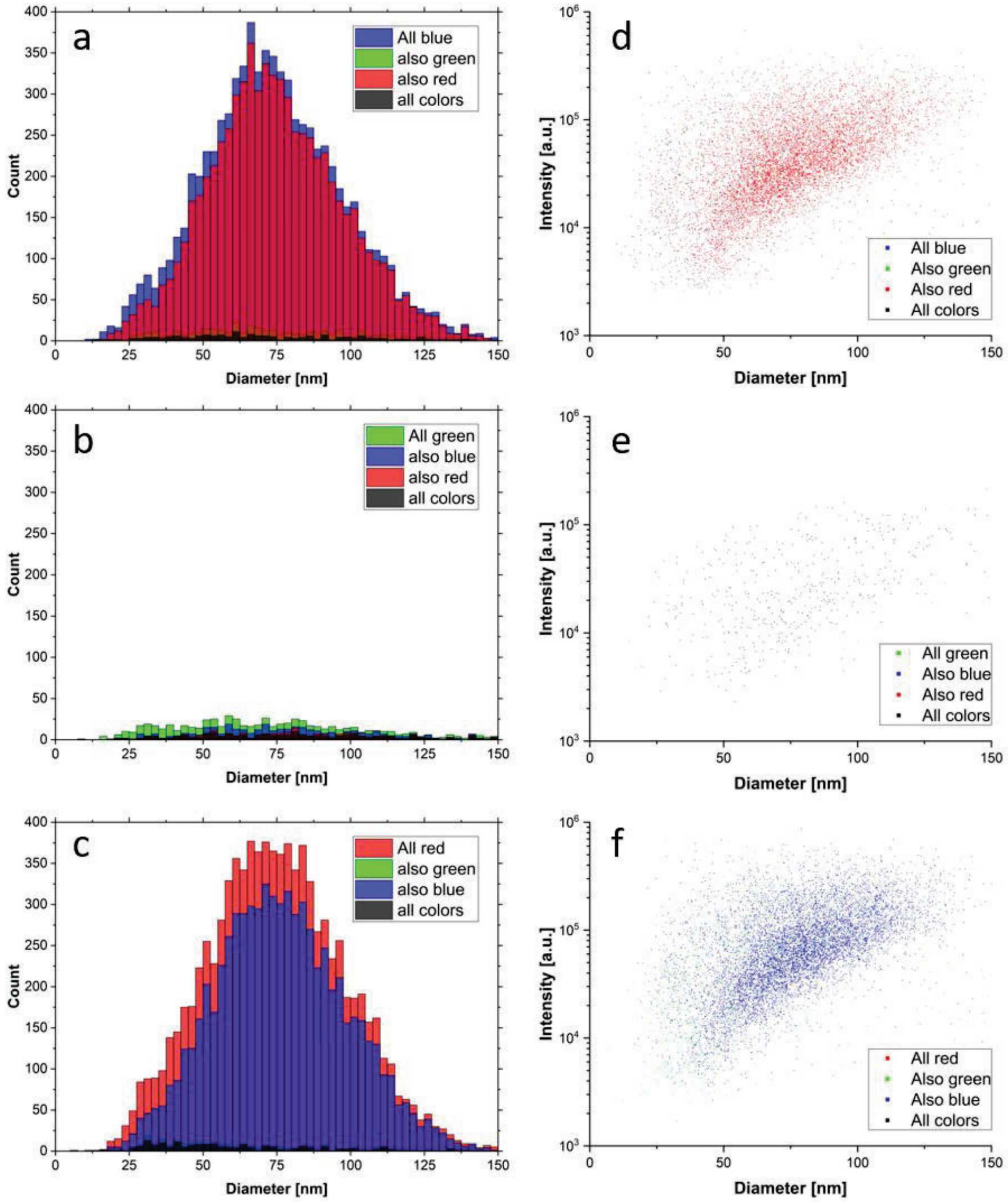

*Supplementary Figure 11. Demonstration of colocalization done with the Nano-SMF for a sample of LUVs POPC/(ATTO488 and ATTO647) with crosstalk correction. To the left are size histograms from the three color channels: a) blue (488 nm), b) green (550 nm) and c) red (647 nm). In the middle are scatter plots of intensity vs diameter from color channel: d) blue (488 nm), e) green (550 nm) and f) red (647 nm).*

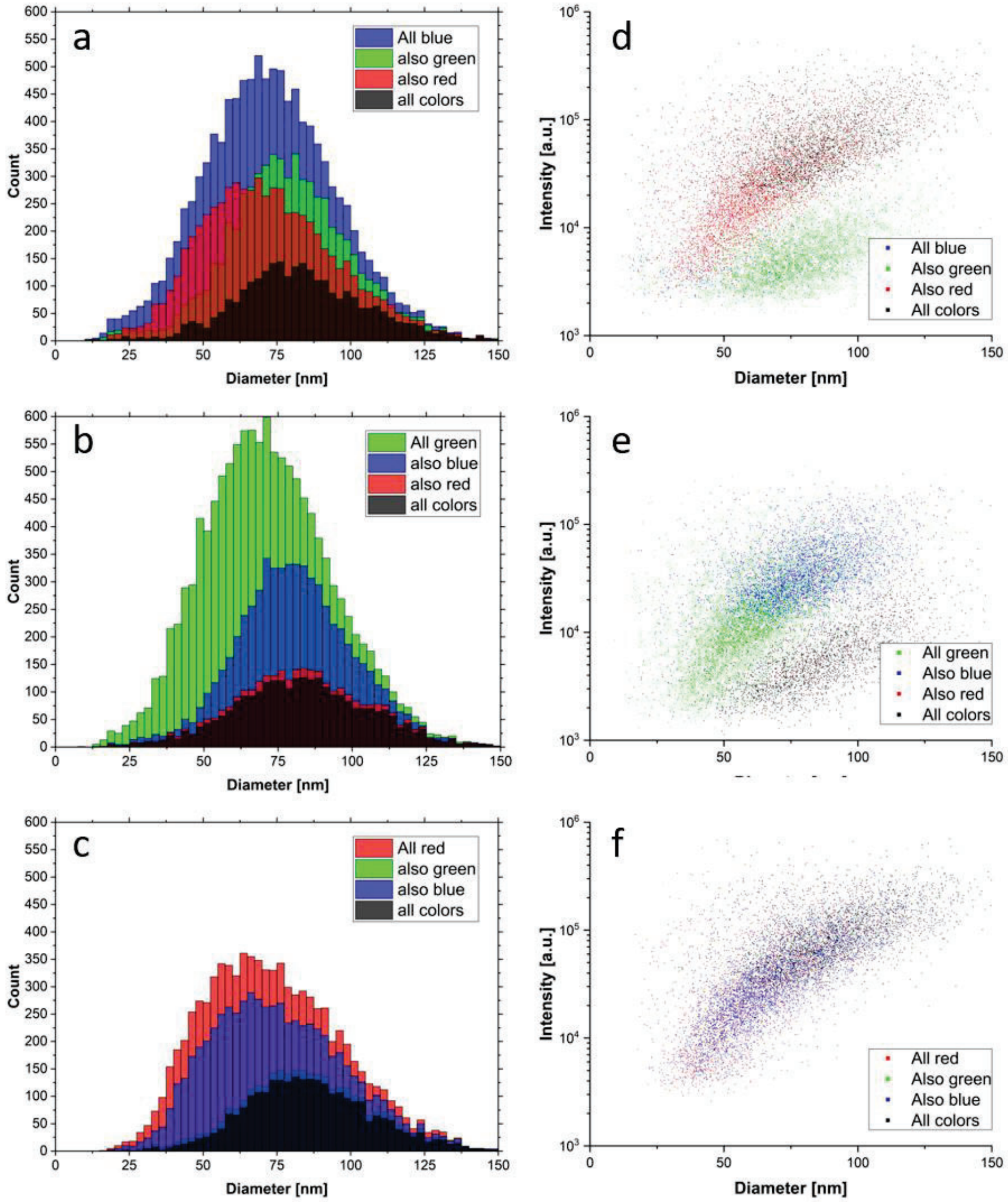

*Supplementary Figure 12. Demonstration of colocalization done with the Nano-SMF for a sample of LUVs POPC/(ATTO488 and ATTO647) and LUVs POPC/Rhod550 without crosstalk correction. To the left are size histograms from the three color channels: a) blue (488 nm), b) green (550 nm) and c) red (647 nm). In the middle are scatter plots of intensity vs diameter from color channel: d) blue (488 nm), e) green (550 nm) and f) red (647 nm).*

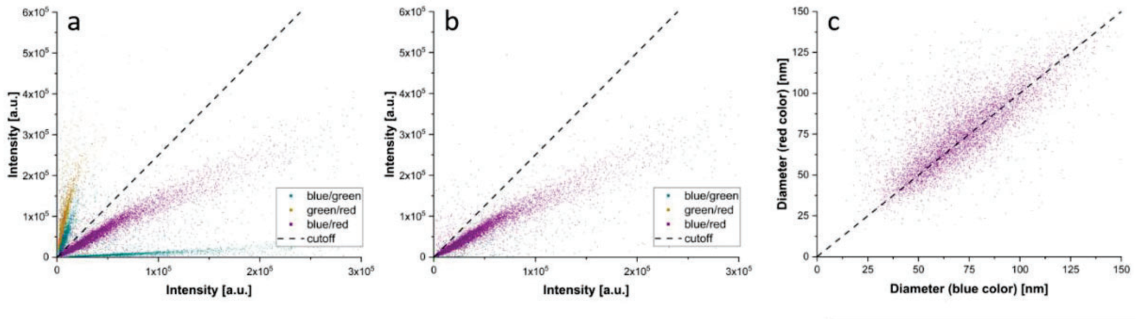

*Supplementary Figure 13. Additional characterization for the colocalization of LUVs POPC/(ATTO488 and ATTO647) and LUVs POPC/Rhod550. a) Scatter plot of intensities from one color vs another. Crosstalk is visible in the form of steep gradient in correlation between the blue/green and the green/red color channels. b) Same as (a) but corrected for crosstalk by the cutoff 0.4 (dashed black line). c) Comparison between sizes determined in the blue color channel and in the red color channel.*

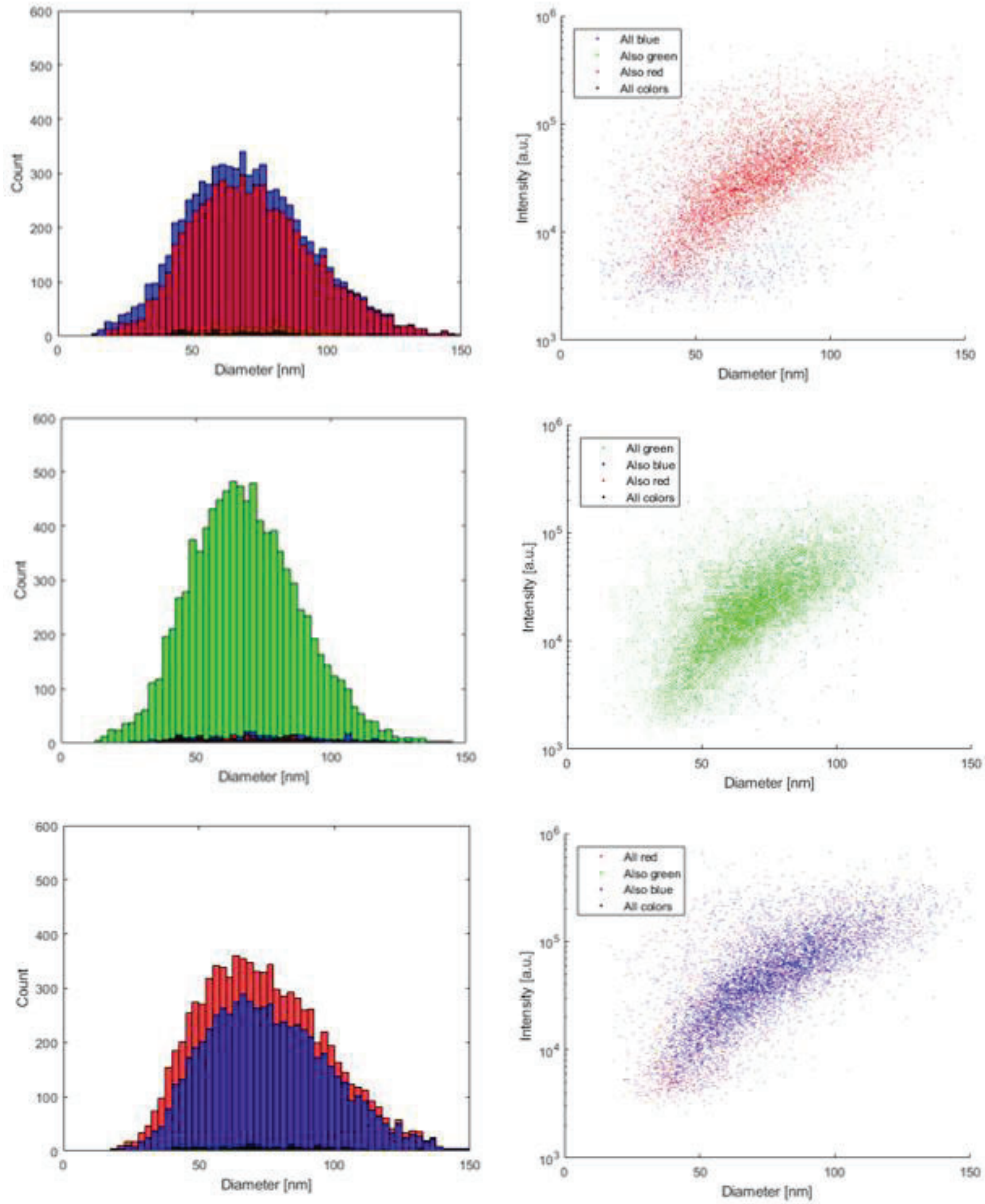

*Supplementary Figure 14. Demonstration of colocalization done with the Nano-SMF for a sample of LUVs POPC/(ATTO488 and ATTO647) and LUVs POPC/Rhod550 with crosstalk correction. To the left are size histograms from the three color channels: a) blue (488), b) green (550) and c) red (647). In the middle are scatter plots of intensity vs diameter from color channel: d) blue (488), e) green (550) and f) red (647).*

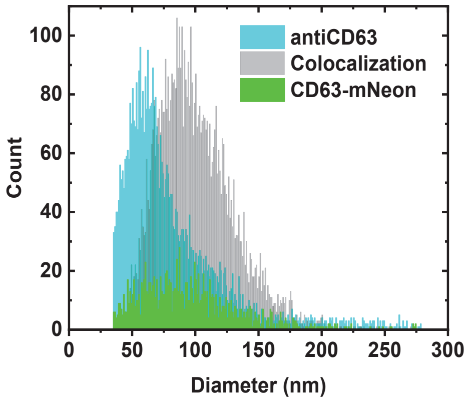

*Supplementary Figure 15. Subpopulation size histogram of CD63-mNeon exosomes (green) labelled with antiCD63 (light blue) characterized with Nano-SMF. Particles colocalizing are represented in grey.*

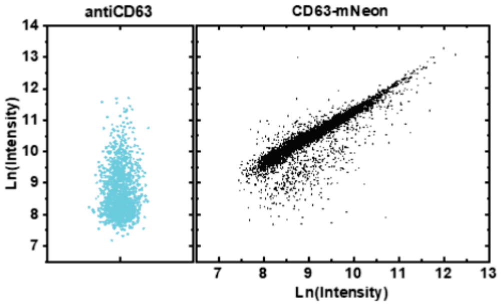

*Supplementary Figure 16. Comparison of fluorescence intensity (antiCD63 channel) versus fluorescence intensity (CD63-mNeon channel) ln-ln plot.*

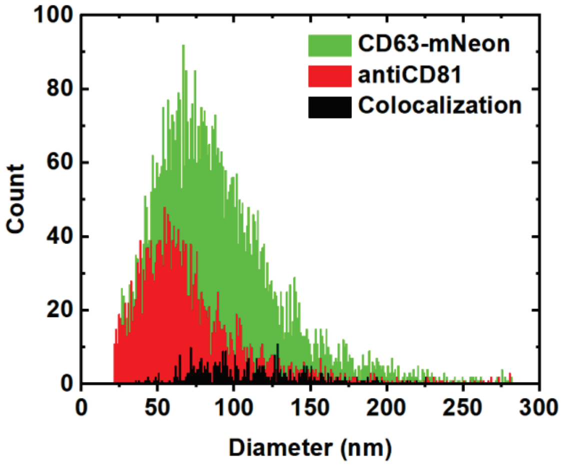

*Supplementary Figure 17. Subpopulation size histogram of CD63-mNeon exosomes (green) labelled with antiCD81 (light blue) characterized with Nano-SMF. Particles colocalizing CD63-mNeon and antiCD81 are represented in black.*

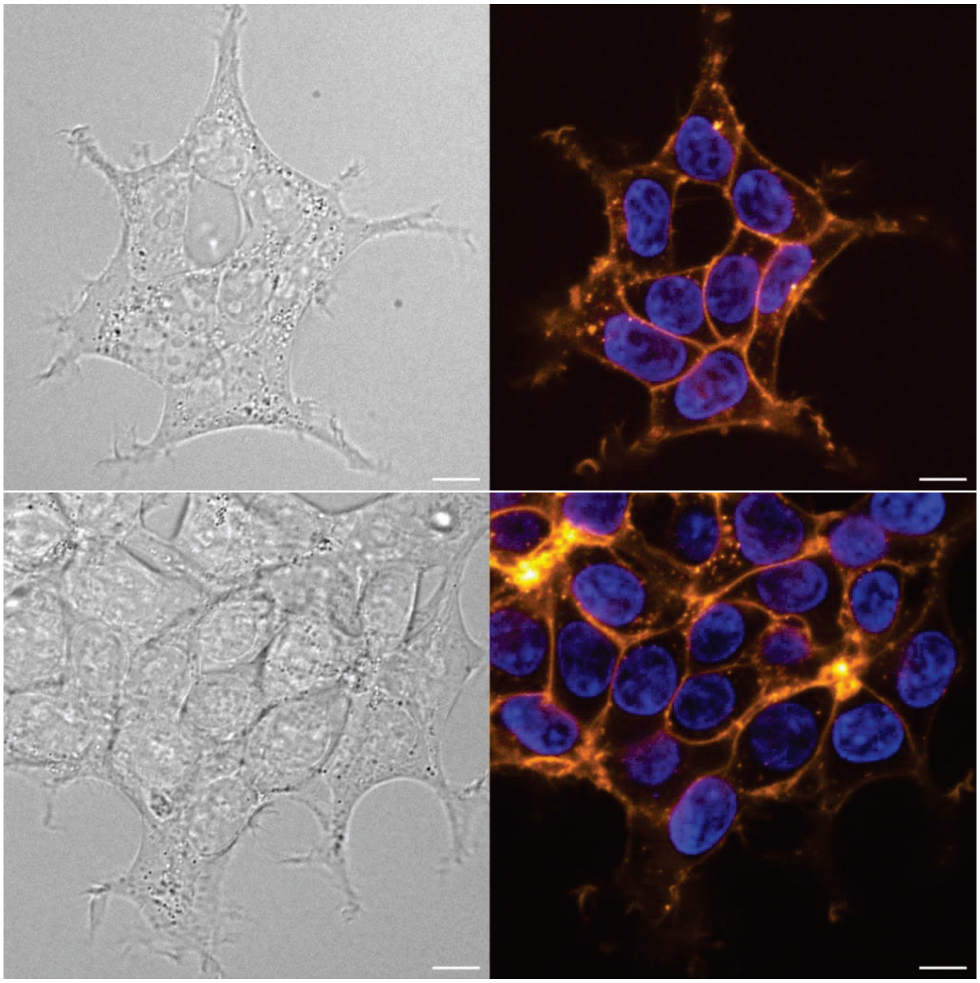

*Supplementary Figure 18. Representative images of HEK293T cells expressing CD63–mCherry acquired using spinning disk confocal microscopy. Bright-field images on the left and a merged fluorescence image on the right, with Hoechst-stained nuclei in blue, and mCherry in orange. The*

*CD63–mCherry signal is displayed using the ImageJ "Orange Hot" lookup table, where increasing intensity is represented from dark orange to white (highest intensity). Scale bar 10 µm.*

### 3. Supplementary Tables

*Supplementary Table 1: Liposome distribution parameters characterized by Nano-SMF and NTA in scattering mode. Mode and Standard Deviation are in nm.*

| Sample | Technique | Median | SE | Mean | SE | Mode | SD | PDI |
|---|---|---|---|---|---|---|---|---|
| POPC/rhod | Nano-SMF | 62.0 | 0.3 | 67.8 | 0.5 | 51.7 | 30.0 | 0.20 |
| | NTA | 81.1 | 0.2 | 85.2 | 0.2 | 73.5 | 27.7 | 0.11 |
| POPC/PKHred | Nano-SMF | 65.7 | 0.1 | 68.1 | 0.2 | 61.3 | 18.5 | 0.07 |
| | NTA | 72.9 | 0.2 | 76.5 | 0.2 | 66.1 | 24.3 | 0.10 |

*Supplementary Table 2: Size and Concentration of liposomes characterized by NTA in scattering or fluorescence mode. Laser wavelength 488 nm. Mode and Standard Deviation are in nm. Concentrations are in particles/mL.*

| | Scattering | | | Fluorescence | | |
|---|---|---|---|---|---|---|
| Sample | Mode | SD | Concentration | Mode | SD | Concentration |
| POPC/ATTO488 | 93.8 | 33.4 | 4.14E12 | 103.2 | 43.3 | 9.55E11 |
| POPC/Rhod550 | 87.2 | 25.9 | 5.70E12 | NA | NA | NA |
| POPC/ATTO647 | 85.4 | 27.3 | 4.81E12 | NA | NA | NA |
| POPC/ATTO488/ATTO647 | 95.0 | 40.8 | 3.18E12 | 108.5 | 32.4 | 1.10E12 |

*Supplementary Table 3: Exosome subpopulation distribution parameters characterized by Nano-SMF. Concentrations are in particles/mL. Median, Mean, Mode, Standard Error and Standard Deviation are in nm. PDI is Polydispersity index.*

| Sample | Label | Nb particles | Concentration | Median | SE | Mean | SE | Mode | SD | PDI |
|---|---|---|---|---|---|---|---|---|---|---|
| Unlabelled | CD63-mNeon | 6227 | 7.7E9 | 81.9 | 0.5 | 89.8 | 0.7 | 78.7 | 40.4 | 0.20 |
| antiCD63 | CD63-mNeon | 1358 | 1.2E9 | 89.2 | 1.2 | 95.5 | 1.1 | 74.2 | 39.4 | 0.17 |
| | antiCD63 | 4609 | 4.0E9 | 66.6 | 0.4 | 82.0 | 0.6 | 58.9 | 41.5 | 0.26 |
| | colocalization | 6147 | 5.3E9 | 96.3 | 0.3 | 100.4 | 0.4 | 87.9 | 29.9 | 0.09 |
| antiCD81 | CD63-mNeon | 6090 | 6.8E9 | 81.9 | 0.5 | 90.5 | 0.5 | 66.6 | 42.9 | 0.22 |
| | antiCD81 | 2506 | 2.8E9 | 63.1 | 0.6 | 69.9 | 0.7 | 51.4 | 33.3 | 0.22 |
| | colocalization | 396 | 4.4E8 | 105.4 | 2.1 | 112.1 | 2.6 | 93.1 | 40.8 | 0.13 |
| Unpurified | CD63-mCherry | 2727 | 2.8E9 | 59.4 | 0.6 | 68.4 | 0.9 | 44.8 | 39.0 | 0.33 |

#### 4. Supplementary video

See shared folder, video is too large.

#### 5. Supplementary References

1 Block, S., Glöckl, G., Weitschies, W. & Helm, C. A. Direct Visualization and Identification of Biofunctionalized Nanoparticles using a Magnetic Atomic Force Microscope. *Nano Letters* **11**, 3587-3592, (2011).

2 Schneider, C. A., Rasband, W. S. & Eliceiri, K. W. NIH Image to ImageJ: 25 years of image analysis. *Nature Methods* **9**, 671-675, (2012).

3 Meijering, E., Dzyubachyk, O. & Smal, I. in *Methods in Enzymology* Vol. 504 (ed P. Michael conn) 183-200 (Academic Press, 2012).

4 Block, S., Fast, B. J., Lundgren, A., Zhdanov, V. P. & Höök, F. Two-dimensional flow nanometry of biological nanoparticles for accurate determination of their size and emission intensity. *Nature Communications* **7**, 12956, (2016).

5 Berglund, A. J. Statistics of camera-based single-particle tracking. *Physical Review E* **82**, 011917, (2010).

6 Edward, J. T. Molecular volumes and the Stokes-Einstein equation. *Journal of Chemical Education* **47**, 261, (1970).

7 Dechadilok, P. & Deen, W. M. Hindrance Factors for Diffusion and Convection in Pores. *Industrial & Engineering Chemistry Research* **45**, 6953-6959, (2006).

8 van der Sman, R. G. M. Drag force on spheres confined on the center line of rectangular microchannels. *Journal of Colloid and Interface Science* **351**, 43-49, (2010).
